# Postmitotic accumulation of histone variant H3.3 in new cortical neurons establishes neuronal chromatin, transcriptome, and identity

**DOI:** 10.1101/2021.10.31.466653

**Authors:** Owen H. Funk, Yaman Qalieh, Daniel Z. Doyle, Mandy M. Lam, Kenneth Y. Kwan

## Abstract

Histone variants, which can be expressed outside of S-phase and deposited DNA synthesis-independently, provide long-term histone replacement in postmitotic cells, including neurons. Beyond replenishment, histone variants also play active roles in gene regulation by modulating chromatin states or enabling nucleosome turnover. Here, we uncover crucial roles for the histone H3 variant H3.3 in neuronal development. We find that newborn cortical excitatory neurons, which have only just completed replication-coupled deposition of canonical H3.1 and H3.2, substantially accumulate H3.3 immediately post mitosis. Co-deletion of H3.3-encoding genes *H3f3a* and *H3f3b* from newly postmitotic neurons abrogates H3.3 accumulation, markedly alters the histone posttranslational modification (PTM) landscape, and causes widespread disruptions to the establishment of the neuronal transcriptome. These changes coincide with developmental phenotypes in neuronal identities and axon projections. Thus, preexisting, replication-dependent histones are insufficient for establishing neuronal chromatin and transcriptome; *de novo* H3.3 is required. Stage-dependent deletion of *H3f3a* and *H3f3b* from (1) cycling neural progenitor cells, (2) neurons immediately post mitosis, or (3) several days later, reveals the first postmitotic days to be a critical window for *de novo* H3.3. After H3.3 accumulation within this developmental window, co-deletion of *H3f3a* and *H3f3b* does not lead to immediate H3.3 loss, but causes progressive H3.3 depletion over several months without widespread transcriptional disruptions or cellular phenotypes. Our study thus uncovers key developmental roles for *de novo* H3.3 in establishing neuronal chromatin, transcriptome, identity, and connectivity immediately post mitosis that are distinct from its role in maintaining total histone H3 levels over the neuronal lifespan.

**Significance:** DNA is packaged around histones into chromatin, which compacts the genome, but also restricts access to DNA. Gene transcription thus requires chromatin reorganization that is precisely regulated, including via variant forms of histones. Here, we find that during a critical developmental window for establishing postmitotic neuronal identity, newly generated cortical excitatory neurons substantially accumulate the histone H3 variant H3.3. Conditional deletion of H3.3-encoding genes from new neurons abrogates *de novo* H3.3 accumulation, and broadly disrupts neuronal histone modifications, gene expression, subtype identity, and axon projections. Thus, preexisting H3 histones are insufficient for establishing neuronal chromatin and transcriptome; *de novo* H3.3 is essential. This developmental requirement for H3.3 is distinct from H3.3 contribution to long-term maintenance of histones in mature neurons.

## Introduction

Chromatin regulation enables dynamic, locus-specific control over gene transcription (1-7). Histone posttranslational modifications (PTMs), chromatin remodeling, and variant forms of histones each contribute to the precise organization of chromatin. Canonical histones are highly expressed in S-phase and assembled into nucleosomes with DNA replication (1, 4, 8, 9); they provide new histones on newly synthesized DNA in dividing cells. Histone variants are expressed throughout the cell cycle and can be incorporated DNA synthesis-independently (1, 4, 9-12); they replenish histones that are turned over in postmitotic cells (10, 13, 14), including terminally postmitotic neurons. In addition to serving as histone replacement, histone variants can also mark specific genomic regions, modulate chromatin states, influence interactions with chromatin regulators, and regulate nucleosome turnover, thereby playing more active roles in specific chromatin functions (2, 4, 15-19).

Histone H3, one of four core histone proteins, is characterized by a protruding N-terminal tail amenable to PTMs that modulate chromatin function, including H3K4me3 and H3K27me3. Canonical H3.1 and H3.2 are encoded by dozens of genes that are clustered, lack introns, and terminate in a stem-loop structure without polyadenylation, which facilitate rapid S-phase transcription (1, 4, 9, 10, 20-24). H3.1 and H3.2 deposition is exclusively DNA synthesis-dependent, relying on histone chaperone CAF-1 for incorporation behind the replication fork. The H3 variant H3.3, in contrast, is encoded by unlinked genes *H3f3a* and *H3f3b*, which undergo splicing, terminate by polyadenylation, and are expressed throughout the cell cycle (1, 4, 9, 10, 20-24). Unlike canonical H3, variant H3.3 can be deposited replication-independently by histone chaperones DAXX or HIRA, preferentially at enhancers, promoters, and gene bodies (15, 17, 25-30), a pattern that suggests H3.3 roles in gene regulation. Functional studies have shown H3.3 involvement in diverse processes, including myogenesis, neural crest differentiation, gametogenesis, genome maintenance, and zygotic genome activation (31-36). In the brain, silencing of H3.3 genes by RNA interference leads to deficits in neuronal layer distribution, activity- and environmental enrichment-induced gene expression, and memory (18, 37).

Here, using conditional genetics to stage-dependently manipulate H3.3-encoding genes *H3f3a* and *H3f3b*, we uncover crucial roles for H3.3 in cerebral cortex development. We find that despite near ubiquitous mRNA expression of *H3f3a* and *H3f3b*, including in cycling cortical neural progenitor cells (NPCs) and postmitotic excitatory neurons, H3.3 protein levels are not uniform. Whereas NPCs are characterized by low H3.3 levels, newborn cortical neurons, which have only just completed replication-coupled deposition of canonical H3.1 and H3.2, undergo substantial accumulation of H3.3 immediately post mitosis. Co-deletion of *H3f3a* and *H3f3b* from newly postmitotic neurons abrogates *de novo* H3.3 accumulation, alters both the deposition and removal of H3 PTMs H3K4me3 and H3K27me3, and causes widespread disruptions to the establishment of the neuronal transcriptome. These transcriptomic changes coincide with neuronal development phenotypes, including in acquisition of distinct subtype identities and formation of axon tracts. Thus, preexisting, replication-dependent histones are insufficient for establishing neuronal chromatin and transcriptome; *de novo*, postmitotic H3.3 is required. Furthermore, by stage-dependent deletion of H3.3-encoding genes from: 1) cycling NPCs; 2) neurons immediately post mitosis; or 3) neurons several days after final mitosis, we delineate the first few postmitotic days to be a critical window for *de novo* H3.3 in cortical neurons. After initial H3.3 accumulation in this developmental window, co-deletion of *H3f3a* and *H3f3b* does not immediately affect H3.3 levels, but causes progressive loss of H3.3 over several months without widespread disruptions to neuronal gene expression, identity, or axon projections. Thus, our study uncovers a developmental role for H3.3 in establishing the chromatin landscape, transcriptome, molecular identity, and axon connectivity in new neurons immediately post mitosis that is distinct from its role in maintaining histone H3 levels over the neuronal lifespan.

## Results

### Substantial postmitotic accumulation of H3.3 in new cortical neurons

H3.3-encoding genes *H3f3a* and *H3f3b* are expressed in many tissues (21). To determine *H3f3a* and *H3f3b* expression in wildtype embryonic day (E)14.5 cortex at the level of single cells, we used single nuclei (sn)RNA-seq. We assayed 9,300 cells that met stringent quality control for snRNA-seq, with a median of 2,642 genes detected per cell. Clustering by Seurat (38) revealed 19 groups that encompassed the full complement of known cell types in embryonic cortex (**Fig. 1A**). Our analysis revealed high *H3f3a* and *H3f3b* expression levels in all types of NPCs, including radial glial cells (RGCs) and intermediate progenitor cells (IPCs), as well as postmitotic neurons at various stages of maturation (**Fig. 1A**). Each of the 19 cell types expressed *H3f3a* and *H3f3b* (**Fig. 1A**), which were detected respectively in 98.0% and 98.2% of cells. Consistent with RNA-seq of developing human cortex (39, 40), our data support that *H3f3a* and *H3f3b* are widely, likely ubiquitously, expressed in cortical development and throughout the cell cycle (21).

**Figure 1.**
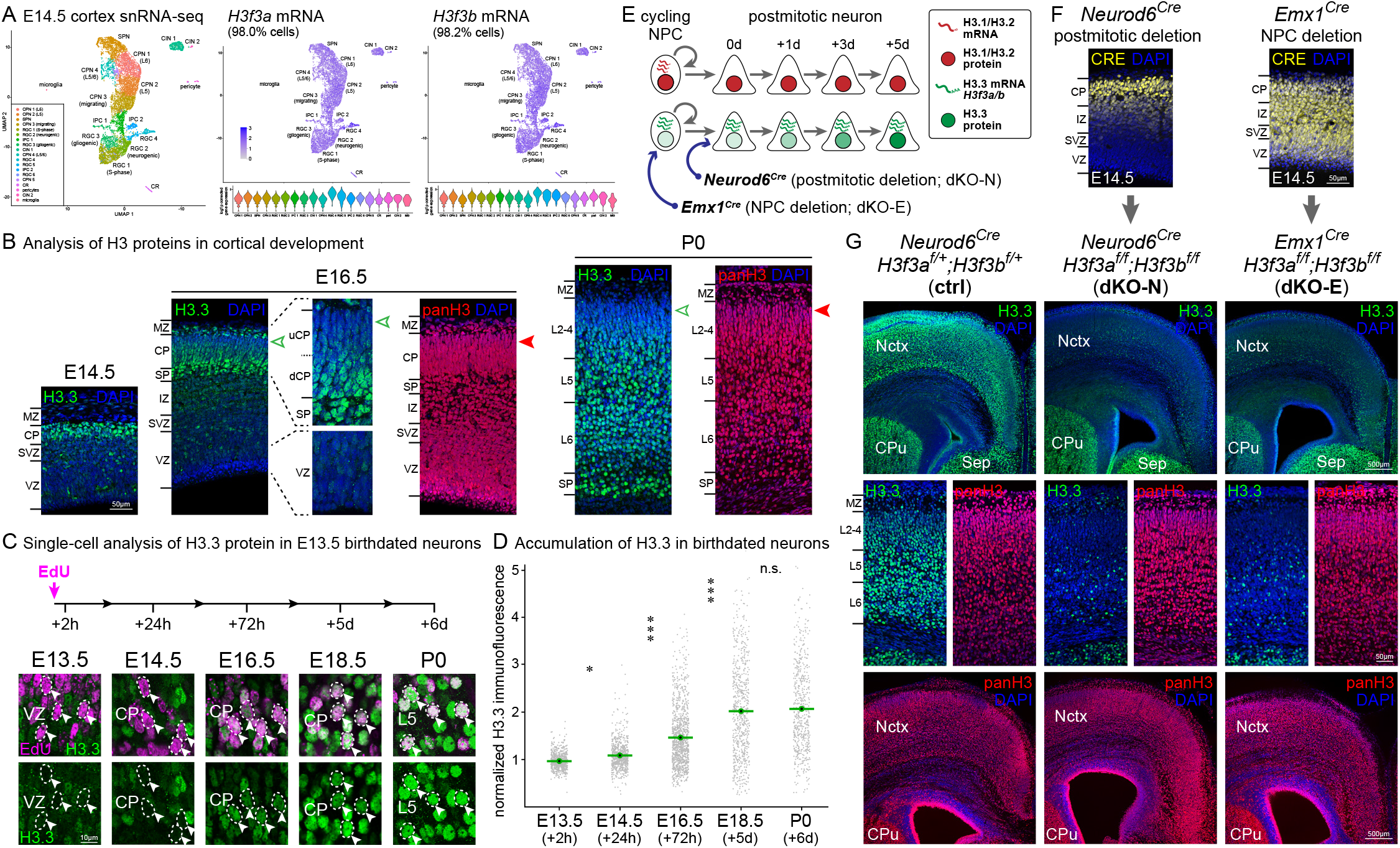
Postmitotic accumulation of H3.3 in cortical neurons and genetic manipulation of *H3f3a* and *H3f3b*. (A) Single nuclei (sn)RNA-seq of wildtype embryonic day (E)14.5 cortex visualized by Uniform Manifold Approximation and Projection (UMAP). *H3f3a* and *H3f3b* mRNA expression was found in each of the 19 identified cell clusters and in 98.0% and 98.2% of cells, respectively. (B) Analysis of H3.3 and pan-H3 protein in cortical development by immunostaining. On E14.5 and E16.5, low levels of H3.3 (green) were present in ventricular zone (VZ) and subventricular zone (SVZ), composed largely of cycling neural progenitor cells (NPCs). High levels of H3.3 were present in cortical plate (CP), composed largely of postmitotic neurons. In E16.5 CP and postnatal day (P)0 cortex, higher levels of H3.3 were present in the early-born neurons of the subplate (SP) and deep layers, compared to the late-born neurons of the upper layers (open arrowheads). Pan-H3 (red) was broadly present and showed no preference for early-born neurons; high levels were present in late-born upper layer neurons (solid arrowhead). (C) Temporal analysis of H3.3 accumulation. Neurons born on E13.5 were labeled by a single pulse of 5-ethynyl-2’-deoxyuridine (EdU). EdU-labeled neurons (magenta, arrowheads) were analyzed by H3.3 immunostaining (green). (D) Quantitative analysis of immunofluorescence revealed a progressive increase in H3.3 levels over the first postmitotic days (one-way ANOVA with Tukey’s post-hoc test, n.s., not significant; ✽, *p* < 0.01; ✽✽✽, *p* < 0.0001). (E) A schematic of H3.1/H3.2 and H3.3 accumulation, and genetic manipulation of H3.3-encoding genes *H3f3a* and *H3f3b*. (F) Analysis of CRE expression (yellow) by immunostaining confirmed the published specificities of *Neurod6*^*Cre*^ in newly postmitotic neurons and of *Emx1*^*Cre*^ in NPCs at E14.5. (G) Analysis of H3 proteins in P0 control (ctrl) and conditional H3.3 genes double knockouts, *Neurod6*^*Cre*^*;H3f3a*^*f/f*^;*H3f3b*^*f/f*^ (dKO-N) and *Emx1*^*Cre*^*;H3f3a*^*f/f*^;*H3f3b*^*f/f*^ (dKO-E). In both dKO-N and dKO-E, H3.3 (green) was largely lost from neocortex (Nctx) but preserved in caudate putamen (CPu) and septum (Sep). The levels of pan-H3 (red) were unaffected in dKO-N or dKO-E. CPN, cortical plate neuron; SPN, subplate neuron; RGC, radial glial cell; IPC, intermediate progenitor cell; CIN, cortical interneuron; CR, Cajal-Retzius cell; MZ, marginal zone; uCP, upper cortical plate; dCP, deep cortical plate; IZ, intermediate zone; L*n*, layer *n*

To assess H3.3 protein levels in individual cells, we used H3.3-specific immunostaining. At E14.5, NPCs in ventricular zone (VZ) and subventricular zone (SVZ) were characterized by low levels of H3.3 protein (**Fig. 1B** and **Fig. S1A**). In contrast, postmitotic neurons in cortical plate (CP) were characterized by high H3.3 levels, suggesting a postmitotic increase. At E16.5, H3.3 levels in NPCs remained low. In CP postmitotic neurons, H3.3 levels were comparatively higher and showed a layer-dependent gradient (**Fig. 1B** and **Fig. S1B**). At E16.5, the earliest born SP and deep layer neurons (41) were characterized by the highest levels of H3.3, and the most recently-born neurons had the lowest levels of H3.3, suggesting progressive H3.3 accumulation over several days after cell cycle exit. In contrast, pan-H3 staining was uniform throughout cortex (**Fig. 1B**). The high pan-H3 levels in NPCs, in which H3.3 was low, was consistent with canonical H3.1 and H3.2 deposition in cycling cells. At P0, the latest born cortical neurons in layer (L)2/3 were characterized by low H3.3 staining, but similar pan-H3 staining compared to earlier-born deep layer neurons (**Fig. 1B**). By P7, both deep and upper layer neurons were characterized by high levels of H3.3 (**Fig. S1C**). This suggests that the layer difference in H3.3 levels at E16.5 and P0 (**Fig. 1B**) reflected the age of neurons since final mitosis; by P7, late-born upper layer neurons have had sufficient time to reach H3.3 levels similar to early-born neurons. To directly quantify H3.3 levels from the time of neuronal birth, we labeled wildtype embryos by a pulse of thymidine analog 5-ethynyl-2’-deoxyuridine (EdU) at E13.5, and tracked labeled cells 2h, 24h, 3 days, 5 days, or 6 days later. We found that in S-phase cells labeled by a 2h pulse of EdU, H3.3 was present only at low levels (**Fig. 1C**). Over the next 6 days following neuronal birth, H3.3 levels progressively increased in EdU-labeled neurons (**Fig. 1C** and **D**). Together, our data suggest that although *H3f3a* and *H3f3b* are expressed throughout the cell cycle, H3.3 histone accumulation in neurons significantly increase postmitotically over several days (**Fig. 1E**).

### Loss of *de novo* H3.3 accumulation in neurons following postmitotic deletion of *H3f3a* and *H3f3b*

*H3f3a* and *H3f3b* encode the same H3.3 protein (1) and constitutive co-deletion of both leads to lethality by E6.5 in mice (36). To study H3.3 in cortical development, we used conditional alleles designed with a reporter; deletion of the *H3f3a* floxed allele would turn on Venus expression, whereas deletion of the *H3f3b* floxed allele would turn on Cerulean (**Fig. S1D**) (42). To assess postmitotic H3.3 function, we used *Neurod6*^*Cre*^ (*Nex*^*Cre*^), which mediates recombination in new excitatory neurons immediately after terminal mitosis, without affecting NPCs (43). To assess H3.3 function in NPCs, we used *Emx1*^*Cre*^, which mediates recombination in cortical NPCs starting at E10.5 (44). We confirmed published patterns of CRE expression at E14.5 (**Fig. 1F**) and cell-type specificity of Cre-mediated H3.3 recombination by reporter expression (**Fig. S1E**). The resulting double H3.3 gene conditional knockouts, *Neurod6*^*Cre*^*;H3f3a*^*f/f*^;*H3f3b*^*f/f*^ (dKO-N) and *Emx1*^*Cre*^*;H3f3a*^*f/f*^;*H3f3b*^*f/f*^ (dKO-E) were born alive. Both dKO-N and dKO-E, however, were characterized by early postnatal lethality within several hours of live birth accompanied by absence of milk spot, which suggested an inability to suckle. Mice with one or more wildtype allele of *H3f3a* or *H3f3b* were not affected by lethality; they lived into adulthood and were fertile as adults.

To assess H3.3 accumulation following *H3f3a* and *H3f3b* deletion, we used H3.3 immunostaining. Neuronal (dKO-N) or NPC (dKO-E) *H3f3a*/*H3f3b* deletion led to an extensive loss of H3.3 from P0 cortex (**Fig. 1G**), but not from subcortical brain structures. Pan-H3 levels, in contrast, were unaffected (**Fig. 1G**). Thus, co-deletion of *H3f3a* and *H3f3b*, in dKO-N or dKO-E, abrogated *de novo* H3.3 synthesis and led to loss of H3.3, but not pan-H3, by P0. H3.3 loss in dKO-N and dKO-E was highly similar (**Fig. 1G**), an observation supported by H3.3 immunoblotting (**Fig. S1F**). This suggests that *H3f3a* and *H3f3b* expression from NPCs, which is preserved in dKO-N, is insufficient for H3.3 accumulation in neurons; postmitotic expression of H3.3 genes is required. This finding is consistent with neurons undergoing significant H3.3 accumulation post mitosis (**Fig. 1B**) and supports that *de novo*, neuronal H3.3 expression is necessary for this increase.

Analysis of whole mount P0 brains showed no significant loss of cortical tissue in dKO-N or dKO-E, whereas quantification of cortical thickness revealed a minor reduction (**Fig. S1G-I**). Importantly, both dKO-N and dKO-E were characterized by an abundance of reporter-expressing cells (**Fig. S1J**), indicating that loss of H3.3 did not cause widespread cell loss, unlike some models of chromatin dysregulation (45, 46). This finding was consistent with immunostaining for cleaved caspase 3 (CC3), a marker of apoptosis, and phospho-(p)KAP1 (TRIM28), a marker of DNA double-strand breaks (**Fig. S1K**), which showed that loss of H3.3 in dKO-N or dKO-E did not cause significant cell lethality or DNA damage at P0.

### Defects in the establishment of the neuronal transcriptome following *H3f3a* and *H3f3b* co-deletion

H3.3 can contribute to transcriptional regulation (11, 15-19, 27). To assess potential transcriptional roles of H3.3 in neurons, we studied the transcriptomic consequences of postmitotic *H3f3a*/*H3f3b* co-deletion in dKO-N cortex at P0. We generated sequencing libraries for unique molecular identifier (UMI) RNA-Seq by ClickSeq (47). We barcoded each individual cDNA molecule with a UMI tag, which enabled highly quantitative measurements of expression levels (48) (**Fig. S2A**). Differential gene expression was analyzed using edgeR (49). This revealed significant changes meeting a stringent FDR of < 0.01 in 948 genes in dKO-N compared to littermate control (**Fig. 2A**). Of these differentially regulated genes (DEGs), 579 were significantly downregulated (log_2_FC < 0) and 369 were significantly upregulated (log_2_FC > 0).

**Figure 2.**
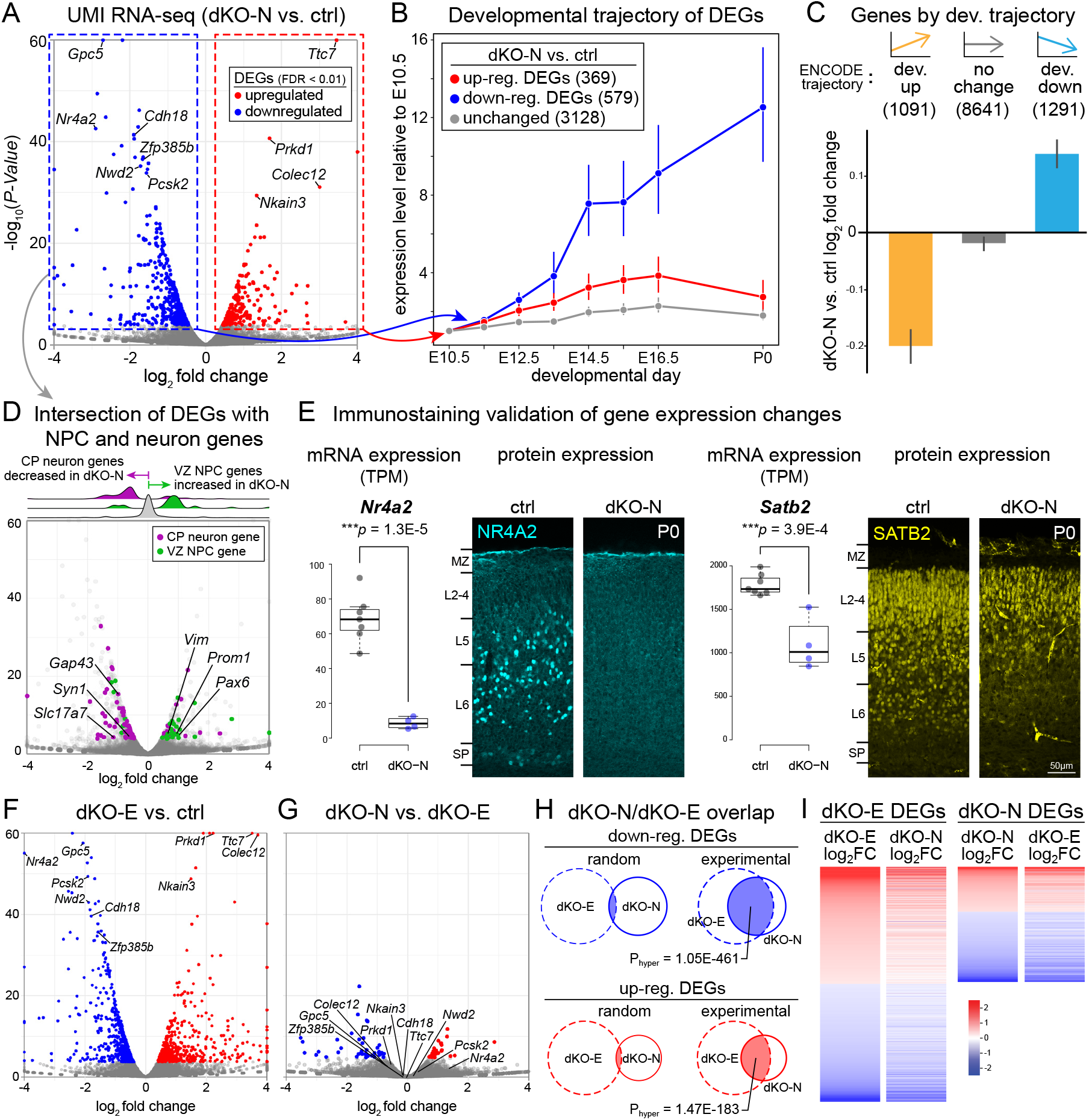
Disrupted developmental acquisition of the neuronal transcriptome following *H3f3a* and *H3f3b* co-deletion. (A) Volcano plot of UMI RNA-seq comparing P0 cortex of dKO-N to control. For each gene, *P* value was calculated with likelihood ratio tests and false discovery rate (FDR) was calculated using the Benjamini-Hochberg procedure. Of 948 differentially expressed genes (DEGs, FDR < 0.01), 579 were downregulated (blue) and 369 were upregulated (red). (B) The developmental expression trajectories of dKO-N DEGs and 3128 unchanged genes (FDR > 0.9, -0.1 < log_2_FC < 0.1, log_2_CPM > 1) based on ENCODE RNA-seq data of wildtype mouse forebrain and normalized to E10.5 levels. Genes downregulated in dKO-N were characterized by a progressive increase in expression across embryonic forebrain development. (C) Genes were categorized based on developmental trajectory using logistic regression on the ENCODE wildtype forebrain RNA-seq data. Genes that normally increase in expression over the course of embryonic development (tangerine, slope > 2, pval < 0.01) were significantly downregulated in dKO-N. (D) Intersectional analysis of DEGs with genes selectively expressed in CP neurons (magenta) or VZ NPCs (green) at E14.5. NPC-selective genes were overrepresented in upregulated DEGs (Phyper = 1.4E-11) and neuron-selective genes were overrepresented in downregulated DEGs (Phyper = 2.8E-43). (E) Immunostaining validation of gene expression changes in P0 cortex. In ctrl, NR4A2 (NURR1, cyan) immunostaining labeled deep layer neurons. In dKO-N, NR4A2 staining was lost. In ctrl, SATB2 (yellow) immunostaining labeled L2-L5 neurons. In dKO-N, SATB2 staining was decreased, especially in the deep layers. (F and G) Volcano plots comparing P0 cortex of dKO-E to control (F) and dKO-N to dKO-E (G). DEGs are indicated as in A. (H) Experimental overlap of dKO-N and dKO-E DEGs compared to random overlap based on hypergeometric probability. (I) Heatmap of dKO-N and dKO-E DEGs based on log_2_FC. dKO-N and dKO-E DEGs showed highly congruent expression change directionality. reg., regulated; dev., developmental; FC, fold change; CPM, counts per million; TPM, transcripts per million

The postmitotic accumulation of H3.3 coincided with an important period of molecular identity acquisition for new neurons (50-61). To determine whether *H3f3a*/*H3f3b* co-deletion affected the developmental establishment of the transcriptional landscape, we intersected genes that were significantly downregulated (579), upregulated (369), or unchanged (3128, FDR > 0.9, -0.1 < log_2_FC < 0.1, log_2_CPM > 1) in dKO-N with ENCODE wildtype forebrain expression data from E10.5 to P0 (62). This revealed in downregulated DEGs an overrepresentation of genes with a positive developmental trajectory (i.e., progressive increase in expression across development) (**Fig. 2B**). This enrichment suggested that following neuronal *H3f3a*/*H3f3b* co-deletion, some genes that normally increase in expression during development failed to do so. We confirmed this finding by using logistic regression on the ENCODE data to identify all genes that normally increase over the course of embryonic forebrain development (i.e., significantly positive trajectory; slope > 2, pval < 0.01) (**Fig. 2C**). As a group, these genes were downregulated in dKO-N, consistent with a role for H3.3 in upregulating neuronal genes to establish the transcriptome post mitosis. Genes that normally decrease over development (i.e., with a significantly negative trajectory; slope < -2, pval < 0.01) were moderately increased in dKO-N, whereas genes that maintained expression levels across development were broadly unaffected (**Fig. 2C**). The transition from cycling NPCs to postmitotic neurons is stark and requires rapid silencing of NPC genes and activation of neuronal genes. To assess whether this transcriptional switch was affected, we intersected DEGs in dKO-N with NPC genes normally enriched in E14.5 VZ and neuronal genes normally enriched in CP (63) (**Fig. 2D**). Our analysis revealed a significant overrepresentation of NPC-selective genes in upregulated DEGs (Phyper = 1.4E-11, 12% of upregulated DEGs), including NPC markers *Pax6, Prom1*, and *Vim*, and a highly significant enrichment of neuron-selective genes in downregulated DEGs (Phyper = 2.8E-43, 16% of downregulated DEGs), including neuronal markers *Syn1, Slc17a7* (*Vglut1*), and *Gap43*. Thus, NPC genes were not appropriately silenced and neuronal genes were not successfully activated in dKO-N. Of the significantly downregulated genes, several are transcription factors that regulate layer-dependent neuronal fates, including *Nr4a2* (*Nurr1*) and *Satb2* (51, 57, 64) (**Fig. 2E** and **Fig. S2B**). We used anti-NR4A2 and anti-SATB2 immunostaining to validate these changes at the protein level in P0 cortex; NR4A2 expression was lost in dKO-N, whereas SATB2 levels were decreased, especially in the deep layers (**Fig. 2E**). Several other neuronal fate markers were also disrupted in dKO-N (**Fig. S2B**). Thus, our results support a requirement for *de novo*, postmitotic H3.3 in silencing NPC genes and activating neuronal genes in postmitotic neurons.

In embryonic and hematopoietic stem cells, H3.3 is required for silencing of endogenous retroviral elements (ERVs), in particular class I and II ERVs (65, 66). Leveraging our transcriptome data, we assessed whether repetitive and transposable elements (67) became derepressed in dKO-N. Interestingly, we found one family of repetitive elements to be significantly upregulated in dKO-N: the class II ERV IAPEY4_I-int|LTR|ERVK (FDR = 5.1E-6, fold change = 1.86, **Fig. S2C**), suggesting that *de novo* H3.3 can contribute to postmitotic silencing of ERVs in neurons. Together, our data support that the broad transcriptomic defects following neuronal *H3f3a*/*H3f3b* deletion include specific deficits in the developmental activation of the neuronal transcriptome and silencing of NPC genes, as well as suppression of some repetitive elements.

To determine whether the comparatively lower levels of H3.3 within NPCs can contribute to gene regulation, we analyzed the P0 cortical transcriptome of dKO-E. This revealed in dKO-E 1934 DEGs, of which 964 were downregulated and 970 were upregulated compared to control (**Fig. 2F**). Although the number of genes meeting FDR < 0.01 was higher in dKO-E compared to dKO-N, analysis of DEGs (**Fig. 2G**) and hypergeometric distributions (**Fig. 2H**) revealed significantly overlapping transcriptomic changes between dKO-E and dKO-N. Furthermore, the genes that were differentially expressed in dKO-E were remarkably consistent in expression change directionality in dKO-N (**Fig. 2I**). Thus, these analyses revealed that *H3f3a*/*H3f3b* deletion from NPCs prior to final mitosis (dKO-E) led to significantly overlapping gene expression changes compared to *H3f3a*/*H3f3b* deletion from neurons postmitosis (dKO-N). Our transcriptomic results thus support an important postmitotic role for H3.3 in the developmental establishment the neuronal transcriptome.

### Disrupted histone posttranslational modifications following neuronal *H3f3a* and *H3f3b* co-deletion

Gene regulation is dependent on histone PTMs, including those decorating H3 (68). To determine whether postmitotic *H3f3a*/*H3f3b* co-deletion affected H3 PTMs, we used CUT&Tag (69) to assess the genomic locations of activating mark H3K4me3 and repressive mark H3K27me3 in dKO-N and control cortex at P0. We first benchmarked our control P0 cortex H3K4me3 and H3K27me3 CUT&Tag data against P0 forebrain ChIP-seq data from ENCODE (62), which revealed consistent and extensive overlap (**Fig. S3A**). Using CUT&Tag, we found in P0 dKO-N striking H3K4me3 and H3K27me3 changes, both significant gains and significant losses, in a locus-dependent manner (**Fig. 3A & B**). At a stringent FDR < 0.05, 220 annotated transcriptional start sites (TSSs) significantly gained H3K4me3, whereas 80 TSSs lost this activating mark in dKO-N. Furthermore, 105 TSSs gained H3K27me3, and 170 lost this repressive mark. These widespread chromatin changes in dKO-N suggest that simple addition and removal of PTMs on preexisting, replication-dependent H3 histones are insufficient to build the neuronal chromatin landscape; *de novo*, postmitotic H3.3 is required.

**Figure 3.**
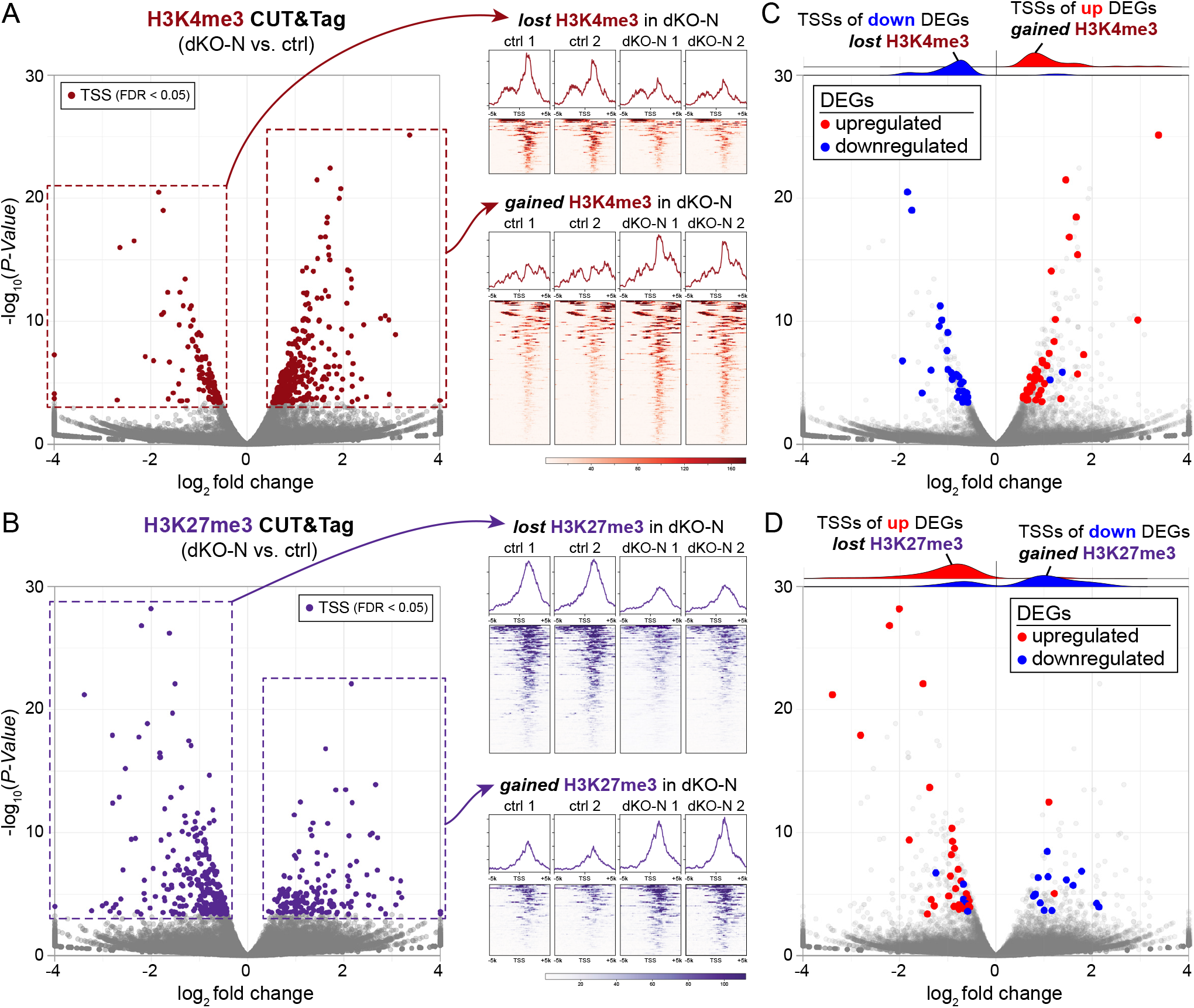
Disrupted histone modifications following *H3f3a* and *H3f3b* co-deletion. (A) Volcano plot of H3K4me3 CUT&Tag signal at annotated transcriptional start sites (TSSs) comparing P0 cortex of dKO-N to control, and normalized H3K4me3 CUT&Tag profiles 5 kilobases upstream and downstream of TSSs. At FDR < 0.05, 300 TSSs showed significantly altered H3K4me3 in dKO-N (burgundy); 220 gained H3K4me3 and 80 lost H3K4me3. (B) Volcano plot of H3K27me3 CUT&Tag signal at TSSs comparing dKO-N to control, and normalized H3K27me3 CUT&Tag profiles. 275 TSSs showed significantly altered H3K27me3 in dKO-N (purple); 105 gained H3K27me3 and 170 lost H3K27me3. (C and D) Intersectional analysis of significantly altered TSSs and DEGs in dKO-N. Downregulated DEGs in dKO-N (blue) largely lost the activating mark H3K4me3 at their TSS but gained the repressive mark H3K27me3. Conversely, upregulated DEGs (red) largely gained the activating mark H3K4me3 at their TSSs but lost the repressive mark H3K27me3. DEGs directionally correlated with PTM changes in dKO-N.

We next assessed whether these marked changes to H3 PTMs could contribute to gene expression changes in dKO-N. Strikingly, we found that downregulated DEGs in dKO-N largely lost the activating mark H3K4me3 at their TSS but gained the repressive mark H3K27me3 (**Fig. 3C & D** and **Fig. S3B & C**). Conversely, the TSSs of upregulated DEGs largely gained H3K4me3 but lost H3K27me3. For example, the TSSs of top upregulated DEGs *Prkd1* and *Ttc7* underwent a change in their valency from repressive H3K27me3 to activating H3K4me3 in dKO-N (**Fig. S3D**). The converse occurred for the TSSs of top downregulated genes *Pcsk2* and *Kcnh1*. Thus, gene expression changes directionally correlated with PTM changes in dKO-N. H3.3 is enriched at bivalent (i.e., H3K4me3 and H3K27me3 co-occupied) chromatin domains (17, 70). To investigate whether disrupted histone modifications in dKO-N can impact bivalent domains, we leveraged ENCODE ChIP-seq data from wildtype E10.5 to P0 forebrain for intersectional analyses (62). We found that the TSSs that significantly gained H3K27me3 in dKO-N are normally characterized by H3K4me3 and H3K27me3 enrichment at earlier timepoints (e.g., E10.5); over the course of development, H3K4me3 is maintained, but H3K27me3 is progressively lost (**Fig. S3E**). The dKO-N gain of H3K27me3 at these loci thus suggested an aberrant retention that could impair the resolution of bivalency over development. Overall, we found that gene expression changes in dKO-N showed a remarkable, directional correlation with PTM changes, strongly suggesting that H3.3 mediates gene regulation, at least in part, via H3 PTMs. Without *de novo* H3.3, regulatory PTMs are broadly and locus-dependently disrupted, and new neurons fail to establish the transcriptome.

To assess whether chromatin accessibility was altered in dKO-N, we carried out ATAC-seq (71) of P0 cortex. Interestingly, whereas TSSs showed robust changes in H3K4me3 and H3K27me3, ATAC-seq revealed in dKO-N only 7 differentially accessible TSSs at FDR < 0.05 (**Fig. S4A**), only one of which overlapped with a DEG. Genome-wide, for all 130,155 ATAC-seq peaks, including those outside of TSSs, the vast majority (>96%) were not differentially accessible in dKO-N (**Fig. S4B**). Only a small minority were differentially accessible (2.1% increased, 1.5% decreased). We intersected these differentially accessible regions (DARs) with ENCODE ChIP-seq data (62) and found moderate overlap with enhancer marks H3K27ac and H3K4me1 (**Fig. S4C**). Overall, postmitotic *H3f3a*/*H3f3b* deletion did not lead to robust changes in genome-wide chromatin accessibility at P0, suggesting that the broad dKO-N transcriptomic changes at this age were unlikely to be the direct result of altered DNA accessibility. Some DARs, however, did overlap with putative enhancer regions that could contribute to gene regulation. Together, our findings uncover crucial roles for *de novo* postmitotic H3.3 in establishing the histone PTM landscape and the transcriptome in new neurons.

### Defects in neuron identities following *H3f3a* and *H3f3b* co-deletion

The cortex is specified into six layers, each with a characteristic composition of molecularly defined neurons (41). To assess laminar development, we first used immunostaining for neuronal marker RBFOX3 (NEUN). In P0 control, RBFOX3 was present in all cortical layers but more intensely labeled L5 and subplate (SP) neurons (**Fig. 4A**). In both dKO-N and dKO-E, RBFOX3 was reduced in all cortical layers, and intense labeling of L5 and SP neurons was lost (**Fig. 4A**). We next used layer markers to assess molecular identities. In control, BCL11B (CTIP2) showed intense staining in L5 neurons and moderate staining in L6 neurons in a stereotyped pattern (53, 58) (**Fig 4B**). In both dKO-N and dKO-E, this distinction was lost; a clear differential staining between L5 and L6 was absent. Furthermore, in control, intense BHLHE22 staining was present in L5 neurons, whereas TLE4 staining was restricted to L6 neurons (**Fig. 4C**). In both dKO-N and dKO-E, the L5 staining of BHLHE22 was lost, whereas TLE4 staining extended beyond L6 and was aberrantly present in L5 (**Fig. 4C**). Neuronal identity is defined by combinatorial gene expression. We thus performed co-immunostaining of BCL11B and TLE4 (**Fig. S5A**), and BCL11B and TBR1 (**Fig. S5B**). In P0 control cortex, L6 neurons were intensely labeled by TLE4 and TBR1 and weakly labeled by BCL11B, whereas L5 neurons were characterized by intense BCL11B labeling and an absence of TLE4 and TBR1, similar to previous reports (58, 59, 72). In dKO-N, L5 and L6 neurons were strongly co-labeled by BCL11B and TLE4 (**Fig. S5A**) and BCL11B and TBR1 (**Fig. S5B**). Thus, in the absence of *de novo* H3.3, deep-layer neurons did not acquire fully distinct L5 versus L6 identities.

**Figure 4.**
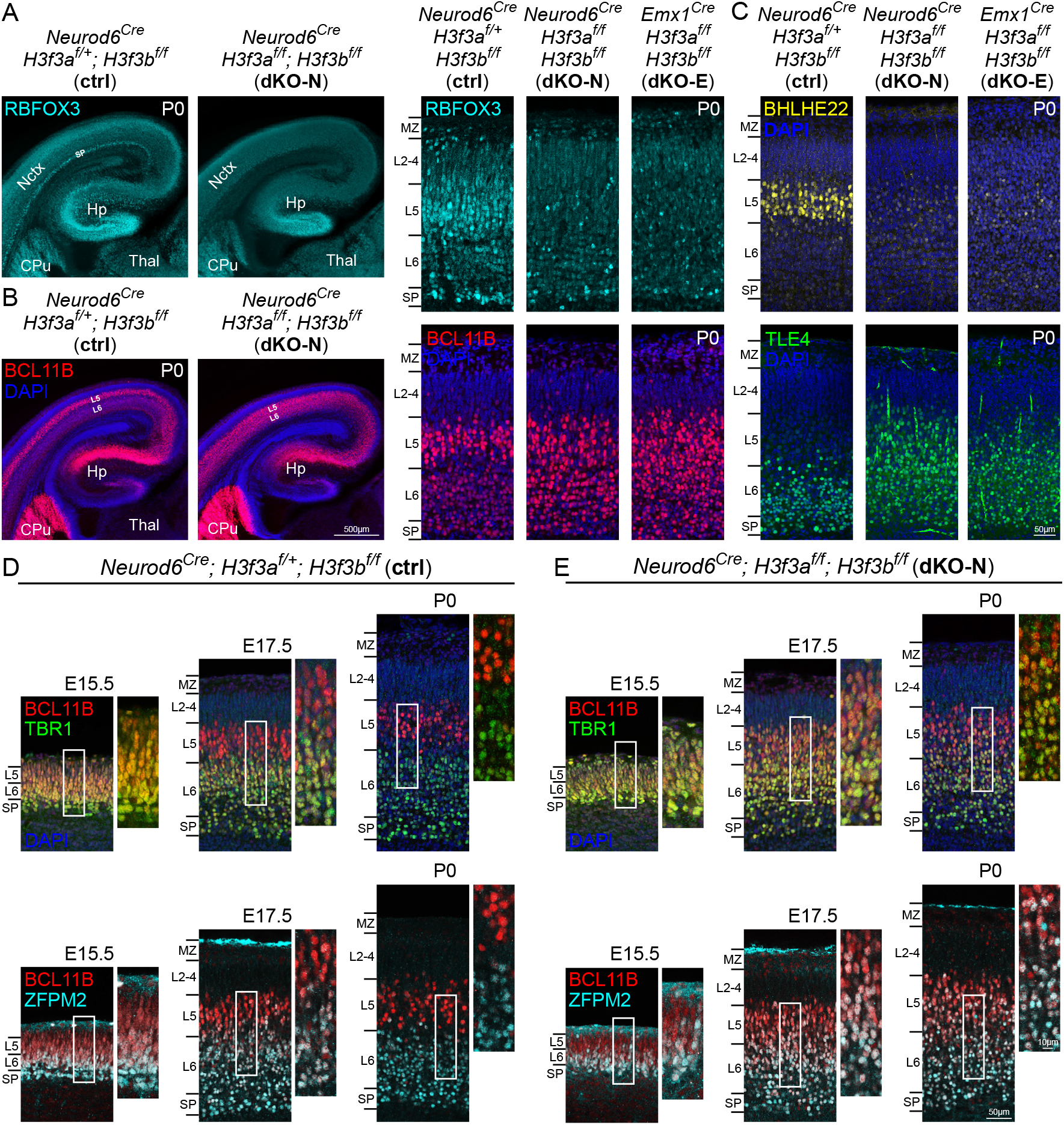
Disrupted establishment of layer-dependent neuronal identities following *H3f3a* and *H3f3b* co-deletion. (A) In P0 control cortex, RBFOX3 (NEUN, cyan) immunostaining was present in all cortical layers and intensely labeled L5 and SP neurons. In dKO-N and dKO-E cortex, RBFOX3 expression was decreased compared to control and intense staining of L5 and SP neurons was absent. (B) In P0 ctrl, BCL11B (CTIP2, red) showed intense staining in L5 neurons and moderate staining in L6 neurons in a stereotyped pattern. In dKO-N and dKO-E, differential BCL11B staining between L5 and L6 was lost. (C) In P0 ctrl, BHLHE22 (yellow) immunostaining intensely labeled L5 neurons, and TLE4 (green) immunostaining was restricted to L6 neurons. In dKO-N and dKO-E, intense L5 staining of BHLHE22 was lost, and TLE4 staining was aberrantly present in L5. (D and E) Postmitotic refinement of deep-layer neuronal identities was analyzed in E15.5, E17.5, and P0 cortex by layer marker co-staining. In control (D), TBR1 (green)- or ZFPM2 (FOG2, cyan)-labeled L6 neurons abundantly co-expressed BCL11B (red) at E15.5. TBR1- or ZFPM2-labeled L6 neurons have begun to downregulate BCL11B at E17.5 and largely did not express high levels of BCL11B by P0. In dKO-N (E), BCL11B was co-expressed with TBR1 or ZFPM2 at E15.5 in a manner similar to control. Abundant marker co-expression, however, persisted at E17.5 and P0. Deep-layer neurons maintained a developmentally immature, mixed L5/L6 molecular identity in dKO-N until P0. Hp, hippocampus; Thal, thalamus

The molecular identities of deep-layer neurons undergo progressive postmitotic refinement (53). For example, newly postmigratory L6 neurons transiently co-express both L5 and L6 markers, but subsequently downregulate L5 markers at late fetal ages to establish their L6-specific identity (53). To determine whether H3.3 contributes to this refinement, we assessed divergence of L5 versus L6 identities over development. As expected, L6 neurons in E15.5 control abundantly co-expressed BCL11B and TBR1, and BCL11B and ZFPM2 (FOG2) (**Fig. 4D**). At E17.5, L6 neurons have begun to downregulate BCL11B, and by P0, TBR1- or ZFPM2-labeled L6 neurons did not express high levels of BCL11B (**Fig. 4D**), consistent with progressive downregulation of L5 expression during acquisition of L6 identity (53). In E15.5 dKO-N, L5 and L6 markers were co-expressed in a manner similar to control (**Fig. 4E**). At E17.5 and P0, however, deep-layer neurons aberrantly maintained abundant L5 and L6 marker co-expression in dKO-N; BCL11B expression had not been downregulated from TBR1- or ZFPM2-labeled L6 neurons. Therefore, in the absence of H3.3, deep-layer neurons maintained mixed L5/L6 molecular identity. To determine whether aberrant marker co-expression was also present in upper-layer neurons in dKO-N, we immunostained for BCL11B and SATB2, mutually repressive markers that respectively label corticofugal and intracortical projection neurons (51, 56) (**Fig. S5C**). In P0 dKO-N, BCL11B and SATB2 labeled largely distinct neurons in the deep layers, and SATB2, but not BCL11B, was present in the upper layers, consistent with control and previous studies (57, 73). Within the deep layers, however, SATB2 expression was reduced (**Fig. 2E**). Together, our analysis of dKO-N revealed a broad spectrum of disruptions to layer-dependent transcription factors, ranging from near complete loss (NR4A2), to shifts in laminar domains (SATB2, TBR1, ZFPM2), to increases in aberrant co-expression (BCL11B with TBR1 or ZFPM2) (**Fig. 2, Fig. 4, Fig. S5**). For deep-layer neurons, we observed both reductions in specific marker expression (NR4A2, SATB2 in deep layers, BHLHE22) and loss of distinct L5 versus L6 identities. For upper layer neurons, we found decreases in marker expression (SATB2, CUX1) without misexpression of deep-layer markers. These changes are consistent with widespread transcriptomic dysregulation in dKO-N and underscore the complex disruptions to cell fate specification following neuronal deletion of *H3f3a*/*H3f3b*.

### Defects in cortical axonal projections following *H3f3a* and *H3f3b* co-deletion

The pathfinding of cortical axons is dependent on correct specification of layer identities (41), and axon guidance mediated by SP neurons (53, 74, 75). We thus analyzed cortical axon tracts (**Fig. 5A**). In P0 control, immunostaining of L1CAM revealed axons traversing white matter (WM) below cortical plate (CP) and corpus callosum (CC) (**Fig. 5B**). *H3f3a*/*H3f3b* co-deletion from neurons (dKO-N) or NPCs (dKO-E) led to a marked loss of white matter thickness, and agenesis of corpus callosum (**Fig. 5B**). Leveraging the Cre-dependent reporters expressed from the *H3f3a* and *H3f3b* floxed loci (**Fig. S1D**), we found in both dKO-N and dKO-E that the anterior commissure (AC) failed to cross the midline and was misrouted toward hypothalamus (**Fig. 5C**). Corticofugal tracts were also disrupted. In both dKO-N and dKO-E, corticofugal axons projected into caudate putamen (CP) and innervated internal capsule (IC) (**Fig. 5D**). Beyond IC, however, corticofugal axons showed deficits in reaching their targets in both dKO-N and dKO-E, including an absence of corticothalamic axons reaching dorsal thalamus (**Fig. 5D**) and an absence of corticospinal tract (CST) axons reaching the pons (**Fig. 5E**). We further analyzed the axons of hippocampal pyramidal neurons (**Fig. S6A**). In both dKO-N and dKO-E, hippocampal axons projected into fimbria, but failed to innervate fornix. These deficits accompanied alterations in hippocampal organization (**Fig. S6B & D**). Together, these findings revealed a requirement for H3.3 in the formation of both intracortical and corticofugal tracts, especially in enabling axons to reach their final targets. These axonal defects were phenotypically indistinguishable between dKO-N and dKO-E, indicating that the requirement for *de novo* H3.3 in axon development is largely postmitotic.

**Figure 5.**
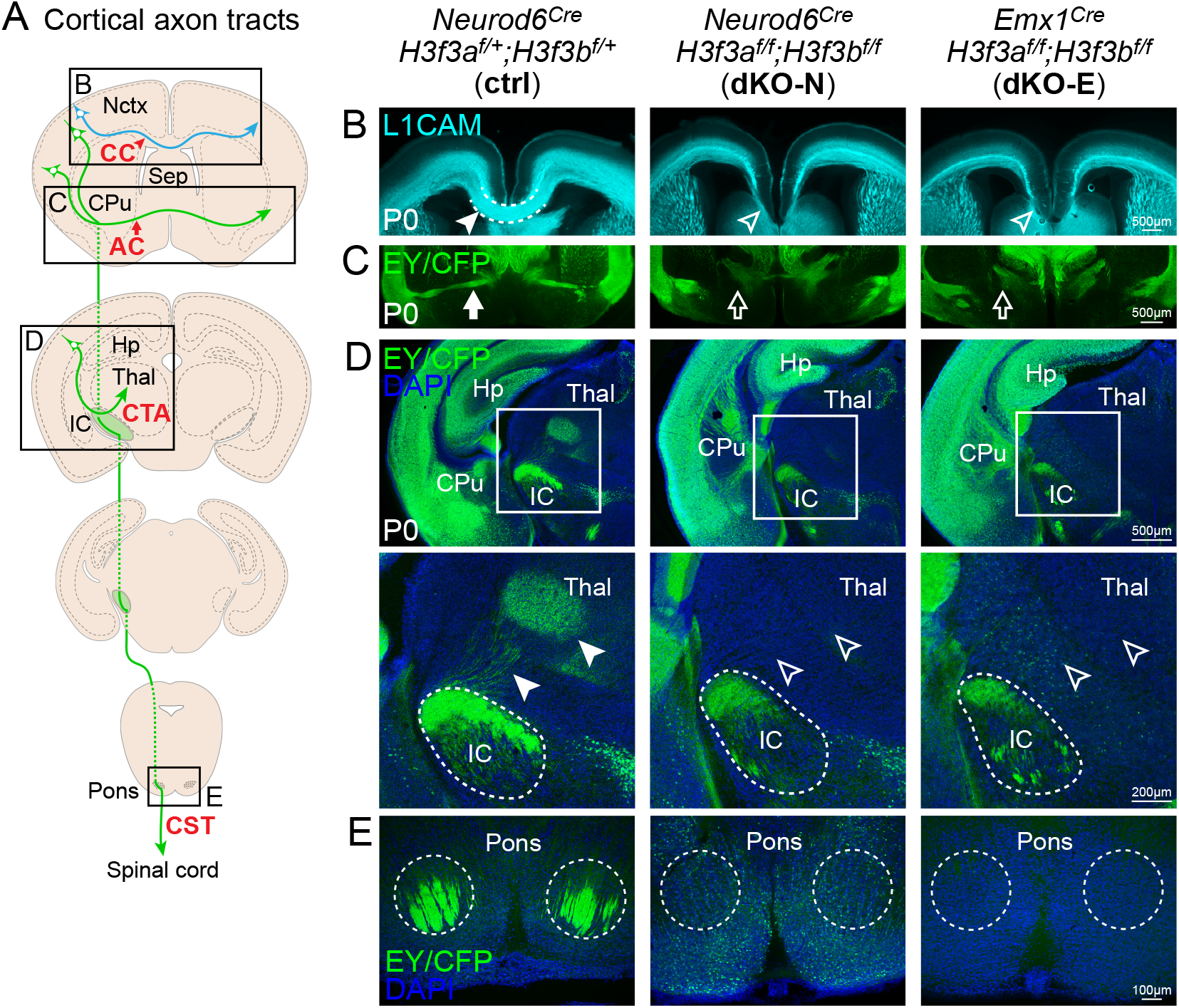
Defective axon tract development following *H3f3a* and *H3f3b* co-deletion. (A) A schematic of major cortical axon tracts. (B) L1CAM (cyan) staining of P0 cortex. dKO-N and dKO-E were characterized by loss of white matter thickness and agenesis of corpus callosum (CC, arrowheads). (C-E) Cre-dependent fluorescent reporters expressed from the *H3f3a* and *H3f3b* floxed loci were detected by anti-EGFP immunostaining of EYFP and ECFP residues. (C) EY/CFP (green) reporter staining revealed failed midline crossing of the anterior commissure (AC, arrows) and misrouting of anterior commissure axons to the hypothalamus in dKO-N and dKO-E. (D) In dKO-N and dKO-E, corticofugal axons reached internal capsule (IC), but corticothalamic tract axons (CTA, arrowheads) failed to innervate thalamus (Thal). (E) Analysis of the corticospinal tract (CST) revealed an absence of axons that reached the level of the pons in dKO-N and dKO-E.

### Co-deletion of *H3f3a* and *H3f3b* after initial H3.3 accumulation

In terminally postmitotic neurons, H3.3 is the primary source of new histone H3 for maintaining overall H3 levels (14, 18). H3.3 is also involved in neuronal senescence, plasticity, and activity-dependent gene expression (18). To assess potential longer term roles of H3.3 in cortical neurons, we generated an additional *H3f3a*/*H3f3b* co-deletion model using Tg(*Rbp4-Cre*) (76). Tg(*Rbp4-Cre*) mediates recombination from L5 neurons starting at ∼E18.5 (**Fig. 6A**), after *de novo* H3.3 accumulation had occurred in the first few postmitotic days. Notably, Tg(*Rbp4-Cre*)*;H3f3a*^*f/f*^;*H3f3b*^*f/f*^ (dKO-R), unlike dKO-N, was not affected by perinatal lethality. Thus, dKO-R (**Fig. 6B**) enabled us to: 1) assess the requirement for *de novo* H3.3 after the initial postmitotic accumulation; and 2) determine the long-term turnover dynamics of H3.3 in mature neurons.

**Figure 6.**
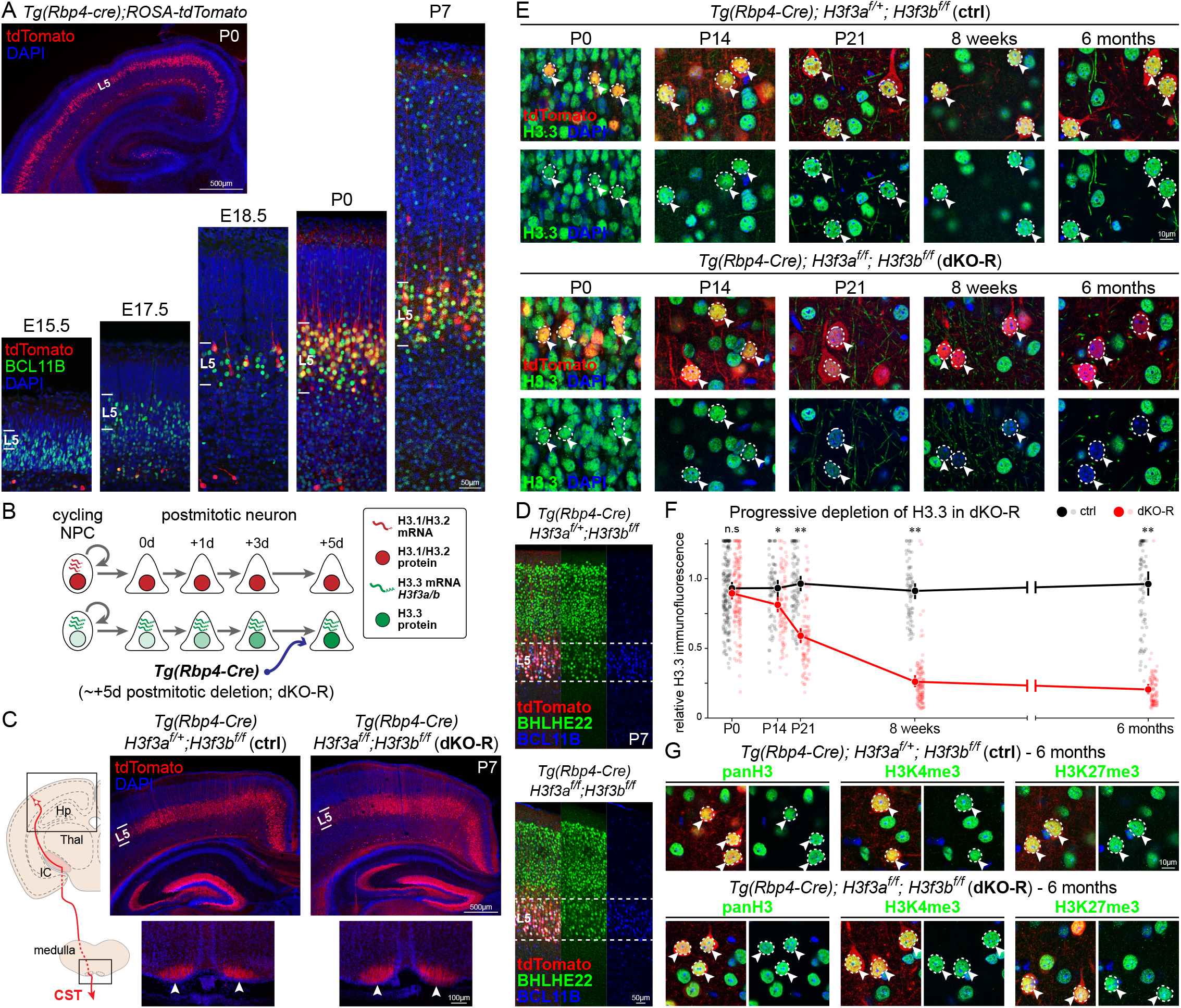
*H3f3a* and *H3f3b* co-deletion after initial H3.3 accumulation in new neurons. (A) Analysis of Tg(*Rbp4-Cre*)-mediated recombination by Cre-dependent reporter *ROSA-tdTomato* (red). Tg(*Rbp4-Cre*)-mediated recombination was present in BCL11B-labeled (green) L5 neurons starting at ∼E18.5, approximately 5 days after terminal mitosis. (B) A schematic of H3.3 accumulation and genetic manipulation of *H3f3a* and *H3f3b* by Tg(*Rbp4-Cre*) in Tg(*Rbp4-Cre*)*;H3f3a*^*f/f*^;*H3f3b*^*f/f*^ (dKO-R). (C) At P7, tdTomato-labeled axons (red) arising from L5 neurons abundantly innervated corticospinal tract (CST) and reached the medullary pyramids (arrowheads) in dKO-R. (D) Analysis of BHLHE22 (green) and BCL11B (blue) by immunostaining revealed normal BHLHE22/BCL11B co-expression in tdTomato-labeled L5 neurons (red) in dKO-R. (E) Temporal analysis of H3.3 levels in tdTomato-labeled L5 neurons (red, arrowheads) by immunostaining. In control cortex, H3.3 (green) is maintained in tdTomato-labeled L5 neurons at each analyzed age. In dKO-R, H3.3 levels were unaffected at P0 but showed a progressive loss from L5 neurons over weeks and months. (F) Quantitative analysis of immunofluorescence (unpaired t-test with Welch’s correction, n.s., not significant; ✽, *p* < 0.01; ✽✽, *p* < 0.001). (G) At 6 months postnatal, pan-H3 and H3 carrying PTMs H3K4me3 or H3K27me3 (green) were assessed in tdTomato-labeled L5 neurons (red, arrowheads) by immunostaining. Despite loss of H3.3, pan-H3 and H3 carrying PTMs were not observably different in dKO-R compared to control.

We first assessed phenotypic differences between *H3f3a*/*H3f3b* co-deletion immediately post mitosis (dKO-N; prior to initial H3.3 accumulation) versus ∼5 days post mitosis (dKO-R; after initial H3.3 accumulation). For direct comparison, we focused on L5 neurons, which were affected by both Tg(*Rbp4-Cre*) and *Neurod6-Cre* deletion. In dKO-N, we found robust axonal phenotypes, including absence of CST axons reaching the pons (**Fig. 6D**). CST axons originate from L5 neurons, which express Tg(*Rbp4-Cre*) (76). We thus used a Cre-dependent tdTomato transgene (*ROSA*^*CAG-tdTomato*^) to label CST axons. In P7 and six-month old dKO-R, we found an abundance of CST axons projecting into and past the pons, and reaching the medullary pyramids, similar to control (**Fig. 6C** and **Fig. S7A**). In dKO-N, we found alterations in layer-dependent neuronal identities; BCL11B staining did not differentiate L5 versus L6 neurons (**Fig. 4B**) and BHLHE22 labeling was lost from L5 (**Fig. 4C**). In P7 dKO-R, however, BCL11B staining differentiated L5 and L6 neurons, and BHLHE22 intensely labeled L5 neurons, each in a manner identical to control (**Fig, 6D**). These data therefore support that after initial H3.3 accumulation, *H3f3a*/*H3f3b* deletion in dKO-R does not disrupt the molecular identity or axon projection of L5 neurons, as *H3f3a*/*H3f3b* deletion prior to initial H3.3 accumulation does in dKO-N.

### Progressive depletion of H3.3 in the absence of *de novo* H3.3

Unlike dKO-N mice, dKO-R mice survived into adulthood, enabling us to examine the long-term dynamics of H3.3. Consistent with substantial H3.3 accumulation in newly postmitotic neurons prior to Tg(*Rbp4-Cre*) expression (**Fig. 1B**), our analysis of P0 dKO-R revealed no immediate change in H3.3 levels in L5 neurons labeled by *ROSA*^*CAG-tdTomato*^ (**Fig. 6E & F**). This contrasted dKO-N, in which *H3f3a*/*H3f3b* co-deletion from new neurons led to a marked loss of H3.3 by P0 (**Fig. 1G**). To determine how H3.3 levels in dKO-R changed over time in the absence of new H3.3, we comprehensively analyzed H3.3 in L5 neurons from P0 to 6 months of age. This revealed in dKO-R a slow, progressive loss of H3.3 from L5 neurons over several months (**Fig. 6E & F**). These data suggested that in cortical neurons, after initial accumulation of H3.3 in the first days postmitosis, H3.3 was turned over much more slowly, over weeks and months. In addition, the abundance of H3.3-negative neurons in dKO-R at 6 months of age indicated that H3.3 was not required for long-term survival of neurons. Consistent with this, our analysis of DNA double strand breaks by pKAP1 marker staining revealed no increase in DNA damage in dKO-R at 6 months compared to control (**Fig. S7B**).

H3.3 is thought to be the key source of H3 throughout the lifespan of postmitotic neurons (14, 18). To assess total H3 levels in dKO-R, we used pan-H3 immunostaining at 6 months, an age at which we observed a 78.7% reduction in H3.3 levels in L5 neurons. Surprisingly, despite marked loss of H3.3, the levels of pan-H3 were not observably different between dKO-R and control (**Fig. 6G**). The presence of histone H3 was confirmed by immunostaining for H3 that carried PTMs H3K4me3 or H3K27me3 (**Fig. 6G**). These data also suggested that H3 PTMs were not eliminated by the lack of *de novo* H3.3 over 6 months. Together, our data showed that following loss of *de novo* H3.3 synthesis, H3.3 levels progressively decreased over several months. Despite loss of H3.3, total H3 levels were maintained up to six months postnatally, suggesting that H3.3 may not be the sole source of H3 proteins in terminally postmitotic neurons.

### Maintenance of neuronal transcriptome following loss of *de novo* H3.3

The initial accumulation of *de novo* H3.3 was required for establishing the transcriptome in new neurons. To assess potential consequences of loss of *de novo* H3.3 on the maintenance of neuronal transcription, we used snRNA-seq to analyze control and dKO-R cortex at 5 weeks, after significant loss of H3.3. Clustering by Seurat (38) revealed 34 groups that encompassed the full complement of known cell types in adult cortex (**Fig. 7A**). Remarkably, the clustering of control and dKO-R cells showed the same clusters in nearly identical UMAP space (**Fig. 7A**) and substantial overlap of each cluster (**Fig. 7B**), including the L5 pyramidal tract (PT) neurons in which Tg(*Rbp4-Cre*) was active (76, 77). This overlap was in striking contrast to the widespread transcriptomic changes we found in dKO-N (**Fig. 2**). We further used differential expression analysis of the L5 PT cluster and specifically assessed whether the DEGs in dKO-N were similarly affected in dKO-R L5 PT. We found that 98.7% of dKO-N DEGs, both up- and down-regulated, became remarkably normalized toward a log_2_FC of 0 (i.e., no change) (**Fig. 7C**). Together, our findings suggest that although the initial accumulation of H3.3 is required to establish the expression of these genes in neurons, once established, *de novo* H3.3 is no longer required to maintain their expression. Our analysis did reveal 71 genes that were modestly differentially regulated in dKO-R L5 PT neurons. Interestingly, a number of these genes are associated with synapse activity, which is consistent with previous reports that in mature neurons, H3.3 can play a longer-term role in neural plasticity (14, 18).

**Figure 7.**
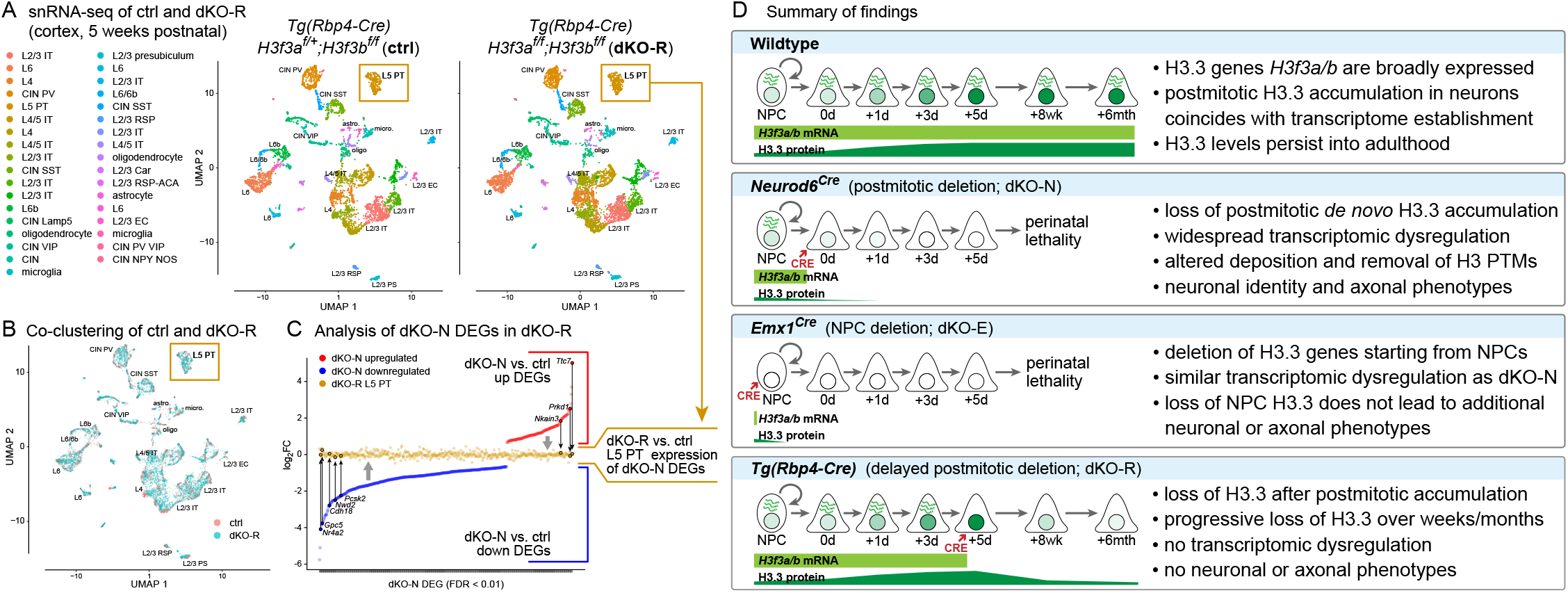
Transcriptome maintenance following *H3f3a* and *H3f3b* co-deletion after initial H3.3 accumulation. (A and B) snRNA-seq of control and dKO-R cortex at postnatal 5 weeks visualized by UMAP. Clustering of ctrl (salmon) and dKO-R (turquoise) cells showed the same 34 cell clusters in UMAP space (A) with substantial overlap of each cluster (B), including the L5 pyramidal tract (PT) neurons (gold boxes) in which Tg(*Rbp4-Cre*) is active. (C) DEGs from dKO-N were compared based on log_2_FC in dKO-N and in dKO-R L5 PT. Both up (red) and down (blue) regulated dKO-N DEGs showed normalization towards log_2_FC of 0 (no change) in dKO-R L5 PT (gold). (D) Summary of key findings. Cortical neurons undergo substantial *de novo* H3.3 accumulation post mitosis. Postmitotic H3.3 is required for developmental establishment of neuronal chromatin and transcriptome, acquisition of distinct neuronal identities, and formation of axon tracts. The first few postmitotic days are a critical window for *de novo* H3.3 in neurons. This early H3.3 accumulation differs in timescale from long-term H3.3 turnover, which occurs over weeks to months.

## Discussion

### Postmitotic H3.3 mediates developmental acquisition of neuronal chromatin and transcriptome

In the first postmitotic days, cortical excitatory neurons undergo precisely regulated molecular and cellular processes to establish their transcriptional identity and axonal connectivity. We find that during this crucial developmental period, newly postmitotic neurons progressively accumulate the histone H3 variant H3.3; this accumulation requires *de novo* H3.3. Our *in vivo* findings are consistent in timing with *in vitro* data from primary neurons, which reach sustained H3.3 levels only after 9 days in culture (18). As a variant histone, H3.3 is thought to serve as the primary source of replacement H3 in terminally postmitotic neurons; in adult brain, H3.3 progressively replaces canonical H3.1/H3.2 over several months after birth (13, 14, 18). In contrast to H3 maintenance over the neuronal lifespan, the late fetal, early postnatal H3.3 accumulation we describe occurs rapidly in new neurons shortly after the final mitosis. This timing is especially notable because canonical H3.1 and H3.2 are produced at high levels and deposited in bulk in dividing cells (22). The requirement for *de novo* H3.3 in newborn neurons, which have only just exited the cell cycle with a full complement of canonical histones, therefore suggests specific, active H3.3 functions distinct from overall H3 maintenance.

Recent studies have identified a number of key transcription factors that function postmitotically to specify the identities of new cortical neurons (50-61). We find that postmitotic co-deletion of *H3f3a* and *H3f3b* (dKO-N) abrogates *de novo* H3.3 accumulation during this critical period, and alters the deposition and removal of H3 PTMs H3K4me3 and H3K27me3, thus disrupting the establishment of the chromatin landscape. Changes to both H3K4me3 and H3K27me3 are consistent with previous reports that H3.3 can mark both active (25, 27) and repressive chromatin (17, 36). An important implication of our findings is that simple addition and removal of PTMs on preexisting, replication-dependent H3 histones are insufficient to build the neuronal chromatin landscape; *de novo*, postmitotic histone H3.3 is required. Without H3.3, regulatory PTMs are broadly and locus-dependently disrupted in dKO-N, and transcription, including that of developmentally regulated genes, is broadly altered. These gene expression changes strongly correlate with PTM changes with the expected directionality. Downregulated genes have lost the activating mark H3K4me3 but gained the repressive mark H3K27me3 in dKO-N. Conversely, upregulated genes have gained H3K4me3 but lost H3K27me3. We further find an expression increase in class II ERVs in dKO-N, consistent with H3.3-dependent repression of ERVs via H3K9me3 (65, 66). Together, our results convergently support the possibility that H3.3 establishes the neuronal transcriptome in large part by mediating establishment of the histone PTM landscape. The requirement for postmitotic H3.3 strongly suggests that the neuronal chromatin landscape is not solely established at the level of NPCs; postmitotic processes play crucial roles. This is consistent with the body of work on transcription factors that specify neuronal fates postmitotically (50-61). We note that although *H3f3a*/*H3f3b* deletion from NPCs (dKO-E) leads to similar transcriptomic changes compared to dKO-N, some changes are unique to dKO-E. Thus, whereas H3.3 plays significant postmitotic roles, the comparatively minor deposition of H3.3 in NPCs may contribute to additional aspects of H3.3-dependent gene regulation.

Histone variants can incorporate into specific genomic regions, interact with chromatin-binding factors, and regulate nucleosome turnover (2, 4, 15-19). Deficits in these mechanisms can contribute to the consequences of *H3f3a*/*H3f3b* deletion in dKO-N. H3.3, which differs from H3.1 and H3.2 at 5 and 4 amino acid residues respectively, is preferentially incorporated into enhancers and promoters (15, 17, 25-30). Thus, intrinsic features of H3.3-containing chromatin can influence these regulatory regions. However, nucleosome core particles containing H3.1/H3.2 versus H3.3 have identical crystal structures (78), and at regulatory regions, nucleosome exchange is equally frequent for H3.1 and H3.3 (29). Of the few amino acid differences, the H3.3S31 residue is absent from H3.1/H3.2. Phosphorylation of H3.3S31 can induce p300-dependent acetylation, particularly at enhancers (79). However, H3.3S31 phosphorylation is carried out by the mitotic checkpoint kinase CHEK1 (79); its potential significance in postmitotic neurons is not known. Incorporation of H3.3 at gene regulatory regions is largely mediated by the HIRA chaperone complex (1, 4, 19). It has been hypothesized that H3.3 can locus-dependently influence chromatin function by recruiting H3.3-specific chaperones, which can in turn nucleate regulatory complexes on H3.3-containing chromatin (9, 79). Whether H3.3 chaperones can locus-dependently recruit a diversity of regulatory complexes to chromatin remains to be elucidated.

Nucleosomes restrict access to genomic DNA. A mechanism that provides and regulates access to DNA is nucleosome turnover, which involves histone removal followed by replacement (10). Nucleosome turnover occurs particularly frequently in regulatory regions (17, 29, 30), and has been proposed to transiently expose regulatory elements to DNA-binding proteins. In addition, nucleosome turnover can shape the chromatin landscape by controlling the valency and chromosomal propagation of histone marks (80), thereby influencing gene regulation in diverse ways. H3.3 plays a crucial role in nucleosome turnover. Knockdown of H3.3 genes significantly stalls nucleosome turnover in neurons and mESCs (10, 18, 19), likely by reducing the soluble pool of H3.3 that is required for nucleosomal assembly and disassembly (10). Thus, stalled nucleosome turnover may contribute to the chromatin and transcriptomic dysregulation in *H3f3a*/*H3f3b* dKO-N. We find that the critical window of H3.3 accumulation coincides with the stark transition from cycling NPCs to postmitotic neurons, and *H3f3a*/*H3f3b* deletion from new neurons impairs the silencing of NPC genes and activation of neuronal genes. H3.3 is known to play diverse roles in cell state transitions, particularly in establishing new identities (31, 34, 37, 79). Immediately after terminal mitosis, nucleosome turnover in new neurons may contribute to removal of PTMs associated with mitotic NPCs and deposition of PTMs appropriate for postmitotic neurons to reset the chromatin landscape. Indeed, in the absence of *de novo* H3.3 in dKO-N, we find locus-dependent gains and losses in H3K4me3 and H3K27me3. Moreover, TSSs with significant H3K27me3 gains in dKO-N include early bivalent loci that normally transition to active chromatin over development. Bivalent domains are enriched for H3.3 in ESCs (17, 70). A potential reduction in nucleosome turnover may affect resolution of bivalency in new neurons, possibly by impairing PTM removal. Nucleosome turnover may also provide access to regulatory elements during establishment of the neuronal transcriptome. Interestingly, our ATAC-seq analysis showed no significant changes in chromatin accessibility at TSSs in dKO-N, despite marked PTM changes. Globally, >96% of ATAC-seq peaks were unaffected in dKO-N. However, among the regions with a significant change in accessibility, some overlap with enhancer marks H3K27ac and H3K4me1. Given the importance of enhancers (81), dysregulation of enhancer function or stalled nucleosome turnover at enhancers may also contribute to the broad transcriptional disruptions in dKO-N.

### Postmitotic H3.3 is required for neuronal and axonal development

The acquisition of neuronal identity occurs under precise regulation, starting with cell fate specification at the level of NPCs (41, 73). Neuronal identities are however not entirely established within NPCs; a number of transcription factors function postmitotically to refine neuronal identities (50-61). Following postmitotic *H3f3a*/*H3f3b* deletion in dKO-N, we find a broad spectrum of disruptions to transcription factors that regulate neuronal fates, ranging from near complete losses, to shifts in laminar domains, to increases in aberrant co-expression. These changes are consistent with widespread transcriptomic dysregulation in dKO-N, and underscore complex disruptions to neuronal fates. Indeed, the broad chromatin perturbations in dKO-N are likely to alter not only the expression of transcription factors, but also their ability activate or repress each other within regulatory circuits. Concomitant with neuronal fate and layer identity phenotypes, we find severe defects in axon development. The P0 dKO-N brain is characterized by agenesis of corpus callosum and misrouting of anterior commissure axons. Corticofugal tracts are able to project into internal capsule, but largely fail to innervate their targets in thalamus or spinal cord. Notably, *H3f3a*/*H3f3b* deletion from NPCs (dKO-E) leads to strikingly similar phenotypes in cortical lamination and axon development compared to dKO-N. This suggests that although NPC deletion of *H3f3a*/*H3f3b* causes some additional transcriptomic changes, these changes do not cause additional phenotypic consequences compared to postmitotic deletion. Thus, the requirement for *de novo* H3.3 in critical molecular and cellular processes of neuronal development is largely postmitotic.

### Long-term dynamics and functions of H3.3 in neurons

*H3f3a* and *H3f3b* are expressed throughout the cell cycle and H3.3 can be deposited DNA synthesis-independently (1, 4, 9-11). In postmitotic neurons, H3.3 is thought to be the primary source of new H3, ultimately replacing canonical H3 genome-wide by several months after birth (13). To assess the requirement for *de novo* H3.3 after the initial postmitotic accumulation, we use Tg(*Rbp4-Cre*) to delete *H3f3a*/*H3f3b* (dKO-R) from cortical L5 neurons ∼5 days after mitosis and subsequent to initial H3.3 accumulation. In phenotypic analyses of dKO-R, we find no disruptions to L5 neuronal identity or axonal projections. The absence of neuronal and axonal phenotypes in dKO-R starkly contrasts dKO-N, and supports that *de novo* H3.3 is required in the first few days after terminal mitosis, but not after initial H3.3 accumulation, for multiple aspects of neuronal development.

As dKO-R is not affected by perinatal lethality, we can explore the long-term dynamics of neuronal H3.3. We find that *H3f3a* and *H3f3b* co-deletion after initial H3.3 accumulation does not immediately affect H3.3 levels. Over the first postnatal months, however, H3.3 levels decrease progressively in the absence of *de novo* H3.3. By 6 months of age, dKO-R L5 neurons have largely lost H3.3 immunostaining. Thus, after initial accumulation of H3.3 in the first postmitotic days, H3.3 is turned over in neurons more slowly, over the course of weeks and months. We next assess total H3 levels in the absence of *de novo* H3.3. We find that at 6 months of age, L5 neurons in dKO-R do not show an observable reduction in overall H3 or in H3 carrying PTMs. Thus, despite loss of H3.3, H3 levels are maintained up to six months postnatally, suggesting that H3 proteins can be produced from a non-H3.3 source in neurons in the absence of *H3f3a*/*H3f3b*. Interestingly, although H3.1 and H3.2 genes are thought to be exclusively expressed in S-phase, there is emerging evidence that under certain scenarios, canonical histone genes can be expressed as polyadenylated transcripts in postmitotic cells (82, 83). It is important to note that although this potential alternate source of H3 may be able to maintain H3 levels over the long term, it is unable to functionally compensate for loss of initial postmitotic H3.3 accumulation in new neurons, as evidenced by the severe transcriptomic and phenotypic consequences in dKO-N.

In the absence of early postmitotic H3.3 accumulation in dKO-N, we find deficits in the developmental acquisition of the neuronal transcriptome. To assess whether loss of *de novo* H3.3 affects transcriptome maintenance, we use snRNA-seq to analyze dKO-R at postnatal five weeks. Remarkably, despite loss of *de novo* H3.3, dKO-R cells formed overlapping clusters with control cells in nearly identical UMAP space. Furthermore, by pseudo-bulk analysis of L5 neurons, we find in dKO-R a remarkable normalization of genes that are differentially expressed in dKO-N. Together, our results suggest that although initial accumulation of H3.3 is required for acquiring the neuronal transcriptome, once established, *de novo* H3.3 is no longer needed to maintain much of the transcriptome. This possibility is consistent with the lack of significant reductions in total H3 or H3 carrying PTMs in dKO-R. Previous studies have reported a role for H3.3 in plasticity-related transcription in adult brain (18). Although we find in dKO-R no changes in transcriptome establishment during development, we do find in mature L5 PT neurons a number of differentially regulated genes that are associated with synaptic activity. Thus, in adult neurons, H3.3 may function by fine-tuning transcription. Together, our study revealed distinct H3.3 functions over the neuronal lifespan. In newly postmitotic neurons, *de novo* H3.3 establishes neuronal chromatin, transcriptome, and identity; in mature neurons, H3.3 provides long-term H3 histones and fine-tunes neuronal transcription.

## Materials and Methods

Further experimental details can be found in SI Appendix, Supplementary Materials and Methods.

### Animals

All animal experiments were performed in compliance with a protocol approved by the University of Michigan Institutional Animal Care & Use Committee.

### ClickSeq

RNA-seq libraries were generated by ClickSeq (47) from 600 ng of purified neocortical RNA.

### CUT&Tag

CUT&Tag was performed from 450k nuclei using the Cut&TagIT Kit following manufacturer’s protocol (Active Motif).

## Acknowledgments

We thank members of the Kwan laboratory, and colleagues in the Michigan Neuroscience Institute (MNI) and Department of Human Genetics for scientific discussions. This work was supported by NIH (R01 NS097525 to K.Y.K., F31 NS110206 to D.Z.D., T32 GM007544 to O.H.F.), Brain Research Foundation (BRFSG-2016-04 to K.Y.K.), March of Dimes Foundation (No. 5-FY15-33 to K.Y.K.), and Simons Foundation Autism Research Initiative (402213, 324586 to K.Y.K.).

