## Supplementary Information for "Postmitotic accumulation of histone variant H3.3 in new cortical neurons establishes neuronal chromatin, transcriptome, and identity"

**This PDF file includes:**

Supplementary Materials and Methods
Figures S1 to S7
SI References

### Supplementary Materials and Methods

#### Experimental Animals

The *H3f3a* (JAX: 021788) and *H3f3b* (JAX: 021789) floxed alleles (1), and *Neuord6*$^{Cre}$ (MGI:2668659) (2), *Emx1*$^{Cre}$ (JAX: 005628) (3), Tg(*Rbp4-Cre*) (MMRRC_031125-UCD) (4, 5), and *ROSA*$^{tdTomato}$ (JAX: 007914) (6) mice were previously generated. For timed pregnancies, the date of vaginal plug was considered embryonic day (E)0.5. Genotyping was performed using DreamTaq Green 2x Master Mix (Thermo Fisher) and genomic DNA isolated from toe or tail clips. Mice were maintained on a standard 12 h day:night cycle with *ad libitum* access to food and water, and all experiments were carried out in compliance with ethical regulations for animal research. Our study protocol was reviewed and approved by the University of Michigan Institutional Animal Care & Use Committee.

Genotyping Primers

| **Gene Target** | **Primer Sequence (5’ -> 3’)** |
|---|---|
| *H3f3a* Mutant Reverse: | CGCGCCCTTCCAAACTAAT |
| *H3f3a* WT Reverse: | TGTTTCCTGGGTGCTTTACC |
| *H3f3a* Common Forward: | GGTGTAATTGAACAGGGAGTGG |
| *H3f3b* Mutant Reverse: | GGTAAAATCCGATAAGGATCGATAG |
| *H3f3b* WT Reverse: | CCTTGAACGTCGCTTGTCTC |
| *H3f3b* Common Forward: | TCTCCCTCACCAATCTCTGG |
| *Cre* Forward: | TCGATGCAACGAGTGATGAG |
| *Cre* Reverse: | TTCGGCTATACGTAACAGGG |

**Immunostaining and Imaging**

Brains were isolated and fixed in 4% PFA overnight at 4°C with agitation and embedded in 4% low-melting agarose for sectioning. Brains were vibratome-sectioned at 70 µm using a Leica VT1000S or VT1200S. Free-floating sections were blocked and immunostained in blocking solution containing 5% donkey serum, 1% BSA, 0.1% glycine, 0.1% lysine, and 0.3% Triton X-100. Free-floating sections were incubated with primary antibodies overnight at 4°C followed by washing 3 × 5-min in PBS and incubated with the corresponding fluorescent secondary antibodies and DAPI in blocking solution for 1h at RT. Following secondary antibody staining, sections were mounted with VECTASHIELD Antifade Mounting Medium (Vector Laboratories). Images were acquired using either an Olympus SZX16 dissecting scope with Olympus U-HGLGPS fluorescent source and Q-Capture Pro 7 software to operate a Q-imaging Retiga 6000 camera, an Olympus Fluoview FV1000 confocal microscope with FV10-ASW software, or an Olympus Fluoview FV3000 confocal microscope with FV31S-SW software.

Primary and Secondary Antibodies

| **Antibody** | **Company** | **Catalog Number** |
|---|---|---|
| Rabbit monoclonal anti-H3.3 | ABclonal | A13824 |
| Rabbit polyclonal anti-panH3 | Abcam | ab1791 |
| Guinea Pig polyclonal anti-Cre Recombinase | Synaptic Systems | 257 004 |
| Goat polyclonal anti-SOX2 | Santa Cruz Biotechnology | sc-17320 |
| Chicken polyclonal anti-RBFOX3 | Sigma-Aldrich | ABN91 |
| Chicken polyclonal anti-GFP | Abcam | ab13970 |
| Rabbit polyclonal anti-GFP | Invitrogen | A-11122 |
| Rabbit polyclonal anti-GAPDH | Santa Cruz Biotechnology | sc-25778 |
| Rabbit polyclonal anti-Phospho KAP-1 | Bethyl | A300-767A |
| Rabbit polyclonal anti-Cleaved Caspase 3 | Cell Signaling Technology | 966 |
| Goat polyclonal anti-NR4A2 | R&D Systems | AF2156 |
| Mouse polyclonal anti-SATB2 | Abcam | ab51502 |
| Rabbit polyclonal anti-CUX1 | ABclonal Custom Project | Project #AP84029 |
| Rat polyclonal anti-BCL11B | Abcam | ab18465 |
| Rabbit polyclonal anti-BHLHE22 | Sigma-Aldrich | HPA064872 |
| Mouse polyclonal anti-TLE4 | Santa Cruz Biotechnology | sc-365406 |
| Rabbit polyclonal anti-TBR1 | Abcam | ab31940 |
| Goat polyclonal anti-FOG-2 | Santa Cruz Biotechnology | sc-9364 |
| Rat monoclonal anti-L1CAM | Sigma-Aldrich | MAB5272 |
| Rabbit polyclonal anti-PROX1 | Sigma-Aldrich | AB5475 |
| Rabbit polyclonal anti-H3K4me3 | Abcam | ab8580 |
| Rabbit polyclonal anti-H3K4me3 | Active Motif | 39060 |
| Rabbit polyclonal anti-H3K27me3 | Millipore | 04-449 |
| Rabbit polyclonal anti-H3K27me3 | Active Motif | 39157 |
| Donkey anti-Goat IgG (H+L), Alexa Fluor 488 | Jackson ImmunoResearch Labs | 805-545-180 |
| Donkey anti-Rabbit IgG (H+L), Alexa Fluor 488 | Jackson ImmunoResearch Labs | 711-545-152 |
| Donkey anti-Rat IgG (H+L), Alexa Fluor 488 | Jackson ImmunoResearch Labs | 712-545-150 |
| Donkey anti-Chicken IgY (H+L), Alexa Fluor 488 | Jackson ImmunoResearch Labs | 703-545-155 |
| Donkey anti-Mouse IgG (H+L), Alexa Fluor 488 | Jackson ImmunoResearch Labs | 715-545-150 |
| Donkey anti-Goat IgG (H+L), Alexa Fluor Cy3 | Jackson ImmunoResearch Labs | 705-165-147 |
| Donkey anti-Rabbit IgG (H+L), Alexa Fluor Cy3 | Jackson ImmunoResearch Labs | 711-165-152 |
| Donkey anti-Rat IgG (H+L), Alexa Fluor Cy3 | Jackson ImmunoResearch Labs | 712-165-153 |
| Donkey anti-Chicken IgY (H+L), Alexa Fluor Cy3 | Jackson ImmunoResearch Labs | 703-165-155 |
| Donkey anti-Rabbit IgG (H+L), Alexa Fluor 647 | Jackson ImmunoResearch Labs | 711-605-152 |

| | | |
|---|---|---|
| Donkey anti-Chicken IgY (H+L), Alexa Fluor 647 | Jackson ImmunoResearch Labs | 703-605-155 |
| Donkey anti-Rat IgG (H+L), Cy5 | Jackson ImmunoResearch Labs | 712-175-153 |

**EdU Birth-Dating**

EdU was given by intraperitoneal injection at a concentration of 5 μg/g. For EdU staining, sections were permeabilized by incubation for 30 min in 0.5% Triton X-100 in PBS followed by 3 × 5-min washes in PBS. EdU staining solution was made fresh each time, containing 100 mM Tris, 4 mM CuSO4, and 100 mM ascorbic acid diluted in 1× PBS. The fluorescently labeled azide molecule was added last, and the staining cocktail was immediately added to sections for a 30-min incubation at room temperature (final concentrations of each fluorescent molecule: 4 mM AlexaFluor488-Azide [Click Chemistry Tools, 1275-1], 10 mM Cy3-Azide [Lumiprobe, A1330], 16 mM AlexaFluor647-Azide [Click Chemistry Tools, 1299-1]). After incubation, sections were washed 3 × 5 min in PBS. After EdU labeling, standard protocols were followed for immunostaining.

**Western blotting**

Mouse cortex was lysed in 1×SDS buffer, proteins were denatured 3 min at 95°C, separated on 4%–12% bis-Tris Gel (NuPAGE), transferred on nitrocellulose membranes, and blocked with 5% non-fat dry milk in Tris Buffer Saline buffer (TBS). Primary incubation was carried out in TBS plus 5% non-fat dry milk, washed 3 times and incubated with anti-rabbit or mouse Peroxidase antibody (1:10,000, Cell Signaling) 1 hour in TBS, followed by washing and ECL Prime chemiluminescence revelation kit (Sigma).

**RNA Isolation**

Embryos were isolated and immediately submerged in ice-cold PBS. Neocortical tissue was dissected and flash-frozen in a dry ice-ethanol bath and stored at −80° until further processing. Tissue was resuspended in 0.5 mL of Trizol and homogenized using metal beads in a bullet blender. Chloroform was added to the sample, and the aqueous phase was isolated following centrifugation for 15 min at >20,000 g at 4 °C. The sample was transferred to a Zymo Research Zymo-Spin IC column and was processed following the manufacturer’s protocol, including on-column DNA digestion. Pure RNA was eluted in DNase/RNase-free water and quantified using a Qubit fluorometer.

**ClickSeq**

RNA-seq libraries were generated by Click-Seq (7) from 600 ng of purified neocortical RNA. Ribosomal RNA was removed from total RNA using NEBNext rRNA Depletion Kit (NEB). ERCC RNA spike-in was included for library quality assessment (Thermo Fisher). SuperScript II (Invitrogen) was used for reverse transcription with 1:30 5 mM AzdNTP:dNTP and 3′ Genomic Adapter-6N RT primer (GTGACTGGAGTTCAGACGTGTGCTCTTCCGATCTNNNNNN). RNaseH treatment was used to remove RNA template and DNA was purified with DNA Clean and Concentrator Kit (Zymo Research). Azido-terminated cDNA was combined with the click adaptor oligo (/5Hexynyl/NNNNNNNNAGATCGGAAGAGCGTCGTGTAGGGAAAGAGTGTAGATCTCGGTGGTCGC CGTATCATT) and click reaction was catalyzed by addition of ascorbic acid and $Cu^{2+}$, with subsequent purification with DNA Clean and Concentrator Kit. Library amplification was performed using Illumina universal primer (AATGATACGGCGACCACCGAG), Illumina indexing primer (CAAGCAGAAGACGGCATACGAGATNNNNNNGTGACTGGAGTTCAGACGTGT) and the manufacturer’s protocols from the 2× One Taq Hot Start Mastermix (NEB). To enrich for amplification products larger than 200 bp, PCR products were purified using Ampure XP (Beckman) magnetic beads at 1.25× ratio. Libraries were analyzed on TapeStation (Agilent) for appropriate quality and distribution and were sequenced at the University of Michigan sequencing core on the Illumina NovaSeq 6000 platform (PE 150 cycles).

**Single-nucleus RNA sequencing**
Cortices were isolated from embryonic and adult mice, and flash frozen in ethanol and dry ice. Nuclei were then isolated and processed for single-nuclei sequencing in accordance with 10x Genomics recommended protocols.
(https://assets.ctfassets.net/an68im79xiti/2IcRfnCKj9MWM4eYZavFnc/47c6023e2987f63ec83173d62124ee7e/CG00055_Demonstrated_Protocol_Dissociation_Mouse_Neural_Tissue_Rev_C.pdf)
Resulting libraries were sequenced at the University of Michigan sequencing core on the Illumina NovaSeq 6000 platform (PE 150 cycles).

**RNA sequencing data analysis**
RNA-seq data were subject to quality-control check using FastQC v0.11.5 (https://www.bioinformatics.babraham.ac.uk/projects/download.html#fastqc). Adapters were trimmed using cutadapt version 1.13 (http://cutadapt.readthedocs.io/en/stable/guide.html). Processed reads were aligned to GENCODE GRCm38 reference genome with STAR (v2.5.2a) (8) and deduplicated according to UMI using UMI-tools (v0.5.3) (9). Read counts were obtained with htseq-count (v0.6.1p1) with intersection-nonempty mode (10). Differential expression was determined with edgeR (11). The *P* value was calculated with likelihood ratio tests and the adjusted *P* value for multiple tested was calculated using the Benjamini-Hochberg procedure, which controls false discovery rate (FDR). Intersectional analysis was done using ENCODE data from embryonic and P0 mouse forebrain. A logistic regression was applied to expression data from every gene, from E10.5 to P0 for each histone mark. From this data, genes were classified as “up” or “down” based on a p-value of <.01 and a slope of > 2 or < -2. bedGraphs were created and normalized using bedtools (v2.29.2) genomecov with -scale, and bigwigs were created using bedgraphToBigWig from UCSC-tools (v2019.09.10). ChIP-seq, CUT&Tag, ATAC-seq, and RNA-seq intersections were done using deepTools2 (12) computeMatrix and computeHeatmap commands. TSS signal for CUT&Tag and ATAC experiments were calculated be first defining a +/- 1kb window around gene TSSs, and then coverage was calculated using bedtools (2.29.2) genomecov. Differential enrichment in these TSS windows was then calculated using edgeR. HOMER (v4.11) analyzeRepeats.pl a was used to assess repetitive elements present in the RNA-seq data, based on consensus sequences taken from RepeatMasker (13).

**ATAC-seq**
Cortical hemispheres were dissected from postnatal P0 mice and lysed using a dounce homogenizer with 10 strokes of pestle A followed by 10 strokes from pestle B in lysis buffer (10 mM Tris·Cl, pH 7.4, NaCl 10 mM, MgCl2 3 mM, 0.1% v/v NP-40). Nuclei were centrifuged at 500g for 10 min at 4 °C. In total, 50,000 nuclei were isolated, and the Omni-ATAC-seq protocol (14) was performed using transposase from Nextera XT Library Preperation Kit (Illumina). After PCR amplification, libraries were purified with the 1.2 × AMPure Beads. Purified ATACseq libraries were analyzed for quality and nucleosome periodicity using a BioAnalyzer High Sensitivity DNA chip (Agilent Technologies) and quantified using the NEBNext Library Quant Kit for Illumina. Libraries were sequenced on an Illumina NovaSeq S4 flow cell to obtain paired-end 150-bp reads. After trimming adapters using cutadapt (1.18) with -minimum-length of 30, reads were aligned to the mouse reference genome GRCm38 using bwa (0.7.15) with default settings (13). Low-quality, mitochondrial, and duplicate reads were removed using a combination of samtools (1.5) and Picard’s MarkDuplicates program (2.8.1). ATAC-seq peaks were called using macs2 (2.1.2) with the parameters: –nomodel –extsize 200 –shift -100, and blacklisted regions were excluded. A consensus set of peaks was generated using bedtools (2.29.2) merge for each condition, and reads from each sample that fell within this consensus peak set were counted using bedtools multicov.

**CUT&Tag**
Cortices were dissected from P0 mice in ice-cold PBS. Tissue was transferred to a Dounce homogenizer containing 1mL Lysis buffer (10 mM Tris-HCL, pH 7.4, 10 mM NaCl, 3 mM MgCl2, 0.1% Tween-20, 0.1%

IGEPAL, 2% BSA). Samples were dounced with 10 strokes pestle A and 10 strokes pestle B. Lysate was strained through a 40 μm filter two times. 2mL of wash buffer (10 mM Tris-HCL, pH 7.4, 10 mM NaCl, 3 mM MgCl2, 0.1% Tween-20, 2% BSA) was used to wash the filter. CUT&Tag was performed using the Cut&TagIT Assay Kit following manufacturer's protocol (Active Motif cat.# 53160). 450k nuclei were used per CUT&Tag reaction. Rabbit primary antibodies were used at 1ug/reaction (histone H3K27me3 antibody (pAb), Active Motif cat# 39157; histone H3K4me3 antibody (pAb), Active Motif cat# 39060). Libraries were sequenced on an Illumina NovaSeq S4 flow cell to obtain paired-end 150-bp reads. Adapters were first trimmed using cutadapt (v1.13) followed by alignment to GENCODE GRCm38 using bwa (v0.7.15), and deduplicated using Picard MarkDuplicates (v2.8.1). macs2 (v2.1.2) was used to call peaks using options -p 1e5 and –keep-dup all with blacklisted regions excluded. A consensus set of peaks was generated using bedtools (2.29.2) merge for each condition, and reads from each sample that fell within this consensus peak set were counted using bedtools multicov. bigWigs were created by first making scaled bedGraphs using bedtools geneomecov with -scale, and then UCSC-utilities bedGraphToBigWig.

**Single-nuclei data processing**

Single nuclei RNA-seq (snRNA-seq) reads were pseudoaligned and counted using kallisto-bus tools (15) with kallisto v0.46.2 and bustools v0.40.0. The kallisto indexes used were generated from the GENCODE GRCm38.p6 reference genome and Ensembl 98 annotations with the lamanno workflow option to include introns. For the Tg(*Rbp4-Cre*) data, transgenes sequences and annotations were added to the index. Seurat v4.0.3 (16, 17) was used to filter, normalize, cluster, and analyze the data. Regularized negative binomial regression (Seurat::SCTransform) was used to normalize the E14.5 data and log normalization was used for the Tg(*Rbp4-Cre*) data. Seurat was then used to determine clusters with original Louvain algorithm and marker genes were found using a Wilcoxon Rank Sum test.

**Image data analysis**

Age- and section level-matched images of dKO and control mice were processed in ImageJ and Adobe Photoshop. Quantification was performed using ImageJ, Cellpose (v0.6) (18) and the Python package openCV.

**Statistical analysis**

Statistical analyses were performed in GraphPad Prism 8 (GraphPad Software). Values were compared using an unpaired Student's *t* test with Welch's correction, or one-way ANOVA with Tukey's post hoc test.

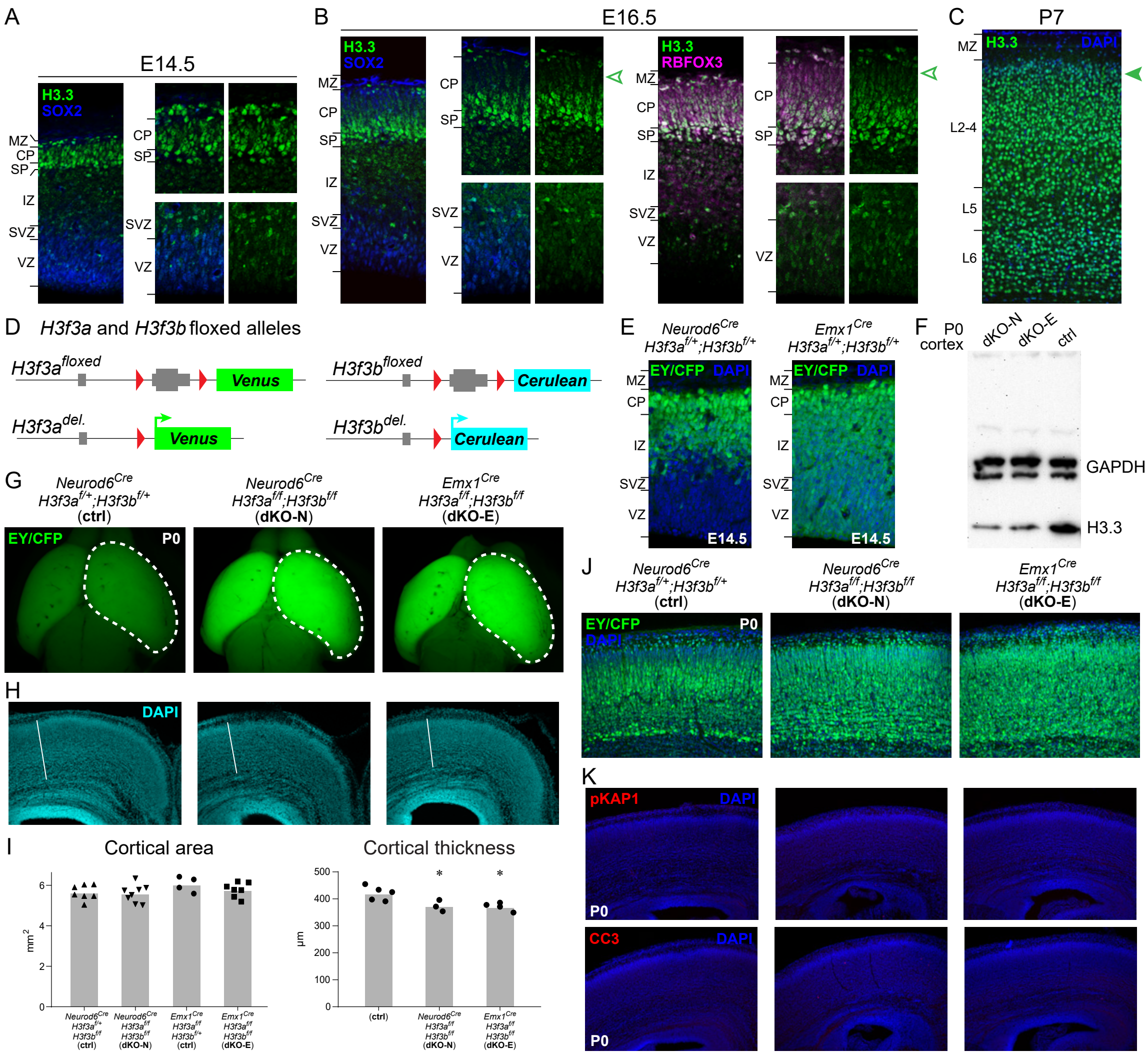


**Fig. S1**

(A and B) Analysis of H3.3 (green) accumulation in developing cortex by immunostaining. (A) On embryonic day (E)14.5, H3.3 levels were low in SOX2+ (blue) NPCs in ventricular zone (VZ) and subventricular zone (SVZ) compared to postmitotic neurons in subplate (SP) and cortical plate (CP). (B) At E16.5, H3.3 levels remained low in SOX2+ VZ and SVZ NPCs. In CP and SP, H3.3 were present at higher levels in RBFOX3+ (magenta) postmitotic neurons and showed a layer-dependent gradient. H3.3 levels were highest in the earliest born SP neurons and deep-layer neurons present in the deep portions of the CP. The most recently-born neurons, present in the uppermost portion of the CP, were characterized by the lowest levels of H3.3 (open arrowheads). This gradient is consistent with progressive H3.3 accumulation over several days in new neurons after their terminal mitosis. (C) H3.3 (green) immunostaining in postnatal day (P)7 cortex. At this age, the late-born upper layer neurons (solid arrowhead) were characterized by high levels of H3.3 similar to early-born deep layer neurons; the gradient present earlier in development had disappeared. (D) A schematic of the *H3f3a* and *H3f3b* floxed alleles. The H3.3 coding sequences were flanked by loxP sites and followed by fluorescent reporter genes *Venus* (*H3f3a*) or *Cerulean* (*H3f3b*). (E) Analysis of reporter gene expression by EGFP immunostaining, which detected EYFP and ECFP residues in Venus and Cerulean (EY/CFP, green). The specificities of *Neurod6*$^{Cre}$ in newly postmitotic neurons and *Emx1*$^{Cre}$ in NPCs were confirmed. (F) H3.3 and GAPDH (loading control) immunoblotting of P0 cortex lysate confirmed similar decreases in H3.3 protein levels in *Neurod6*$^{Cre}$;*H3f3a*$^{f/f}$;*H3f3b*$^{f/f}$ (dKO-N) and *Emx1*$^{Cre}$;*H3f3a*$^{f/f}$;*H3f3b*$^{f/f}$ (dKO-E). (G) Dorsal view of whole mount P0 ctrl, dKO-N, and dKO-E brains. (H) Coronal sections of P0 ctrl, dKO-N, and dKO-E stained by DAPI (cyan). (I) quantitative analysis of cortical hemisphere area and cortical thickness (data are mean, one-way ANOVA with Tukey's post-hoc test; ✻, $p < 0.05$). (J) Analysis of reporter gene expression by EGFP immunostaining (EY/CFP, green) on P0 coronal sections. An abundance of reporter-expressing cells was present in both dKO-N and dKO-E cortex. (K) Immunostaining of DNA double strand break marker pKAP1 (red) and apoptosis marker cleaved caspase 3 (CC3, magenta) revealed no significant DNA damage or cell death in P0 dKO-N or dKO-E.

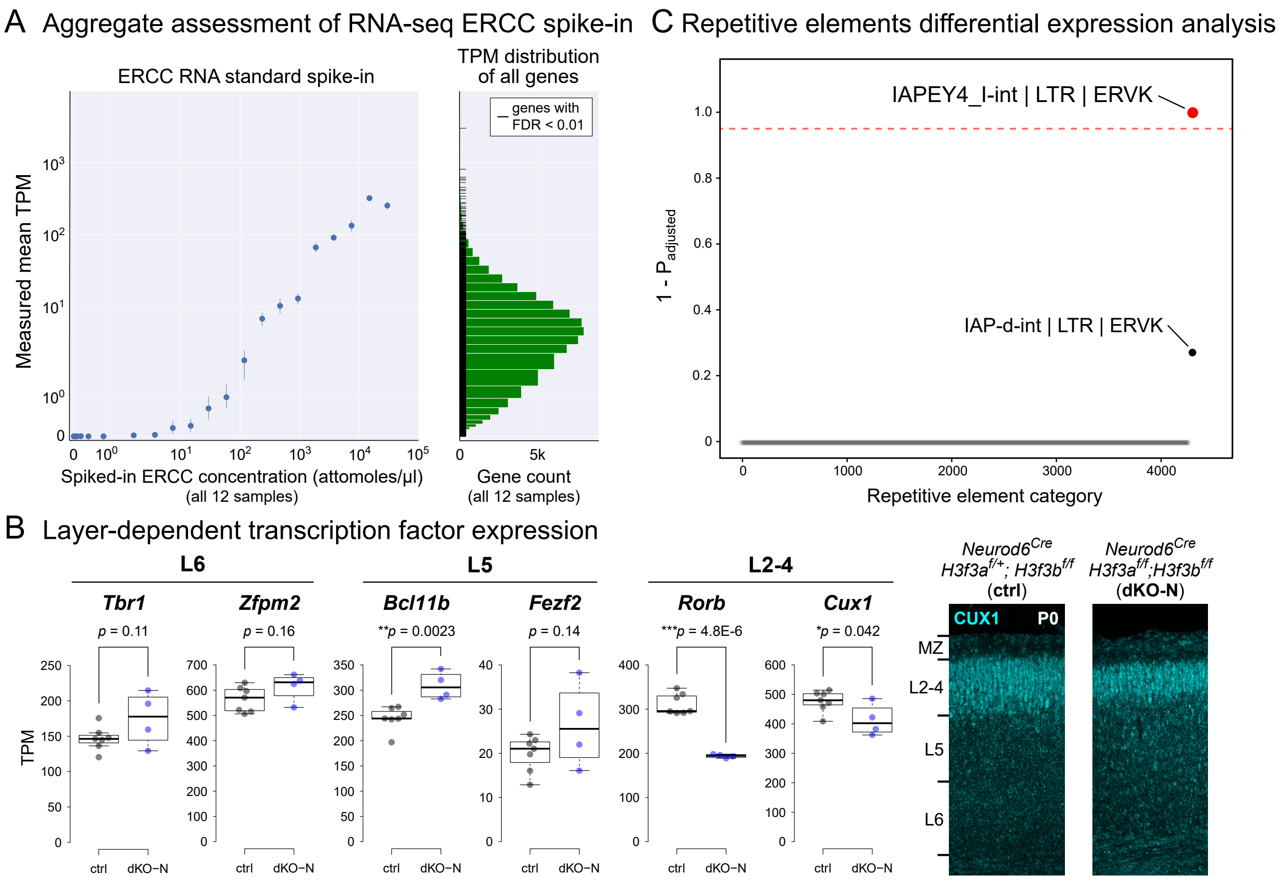


**Fig. S2**

(A) Aggregate assessment of ERCC spike-in standards in UMI RNA-seq revealed excellent quantification over a broad range of expression levels. TPM, transcripts per million. (B) Analysis of layer-dependent transcription factor expression in P0 cortex using RNA-seq data. At the mRNA level, transcription factor expression showed a diversity of disruptions in dKO-N, including reductions in L2-4 markers *Rorb* and *Cux1*. *Cux1* expression changes were validated by immunostaining in P0 cortex. In ctrl, CUX1 (cyan) immunostaining labeled L2-4 neurons. In dKO-N, CUX1 staining was present in the upper layers, but its domain was reduced, consistent with decreased mRNA expression. (C) Analysis of repetitive and transposable elements expression using P0 cortex RNA-seq data. One family of repetitive elements was significantly upregulated in dKO-N: the class II ERV IAPEY4_I-int|LTR|ERVK (FDR = 5.1E-6, fold change = 1.86).

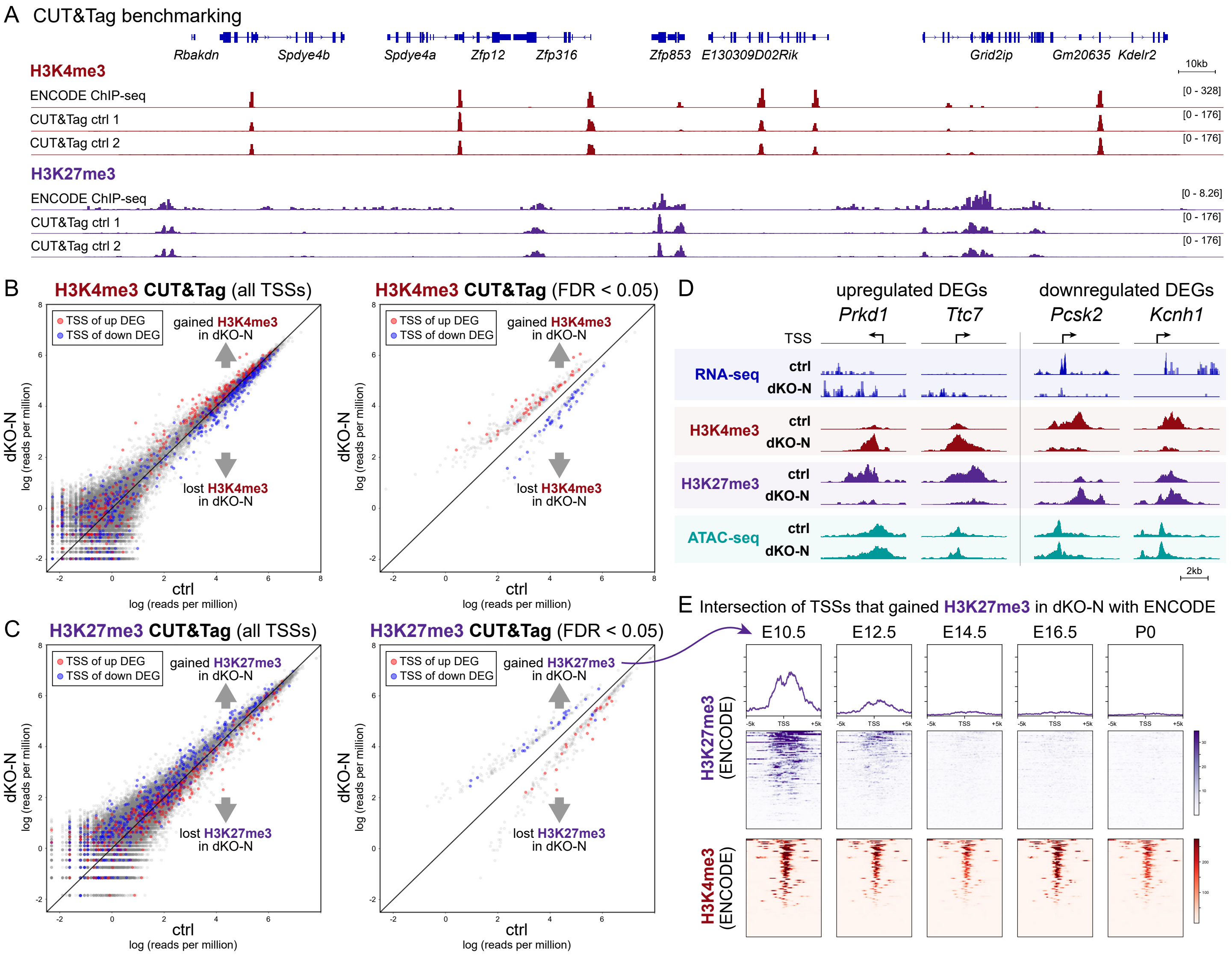


**Fig. S3**

(A) Comparison of P0 control cortex H3K4me3 and H3K27me3 CUT&Tag data against benchmark P0 forebrain ChIP-seq data from ENCODE. Control CUT&Tag data showed extensive and consistent overlap with ENCODE benchmark. (B and C) Scatterplots comparing H3K4me3 (burgundy) or H3K27me3 (purple) CUT&Tag signal at all TSSs or significantly altered TSSs (FDR < 0.05) in dKO-N (y-axis) and ctrl (x-axis). The TSSs of upregulated DEGs (red) and downregulated DEGs (blue) in dKO-N are indicated. Downregulated DEGs largely lost the activating mark H3K4me3 at their TSS but gained the repressive mark H3K27me3. Conversely, upregulated DEGs largely gained H3K4me3 but lost H3K27me3. (D) The TSSs of top upregulated DEGs *Prkd1* and *Ttc7* underwent a change in valency from repressive H3K27me3 to activating H3K4me3 in dKO-N. Conversely, the TSSs of top downregulated genes *Pcsk2* and *Kcnh1* underwent a change from H3K4me3 to H3K27me3 in dKO-N. Chromatin accessibility was not significantly changed at these TSSs. (E) Intersectional analysis of TSSs that gained H3K27me3 in dKO-N with ENCODE forebrain H3K4me3 (burgundy) or H3K27me3 (purple) ChIP-seq data from E10.5 to P0. The TSSs with significant H3K27me3 gains in dKO-N are normally characterized by H3K4me3 and H3K27me3 co-occupancy at E10.5. This bivalency is resolved over the course of development; H3K4me3 enrichment is maintained until P0, but H3K27me3 enrichment is progressively lost.

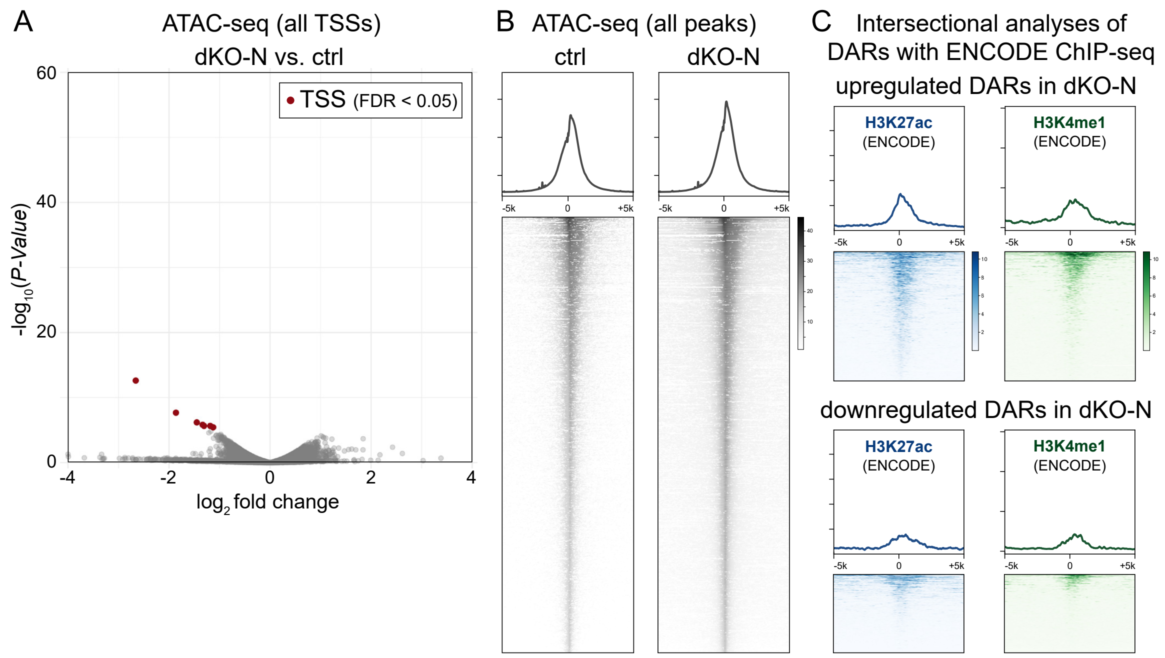


**Fig. S4**

(A) Volcano plot of ATAC-seq signal at annotated TSSs comparing P0 cortex of dKO-N to control. The seven differentially accessible TSSs (FDR < 0.05) are indicated (burgundy). (B) Normalized ATAC-seq profiles 5 kilobases upstream and downstream of all 130,155 ATAC-seq peaks. The vast majority (>96%) of these ATAC-seq peaks were not differentially accessible in dKO-N, suggesting that loss of H3.3 from cortical neurons did not broadly affect chromatin accessibility at P0. (C) Intersectional analysis of differentially accessible regions (DARs) with ENCODE forebrain H3K27ac (dark cerulean) and H3K4me1 (dark green) ChIP-seq data from P0 forebrain. Some DARs coincided with these marks, suggesting a potential overlap with enhancers.

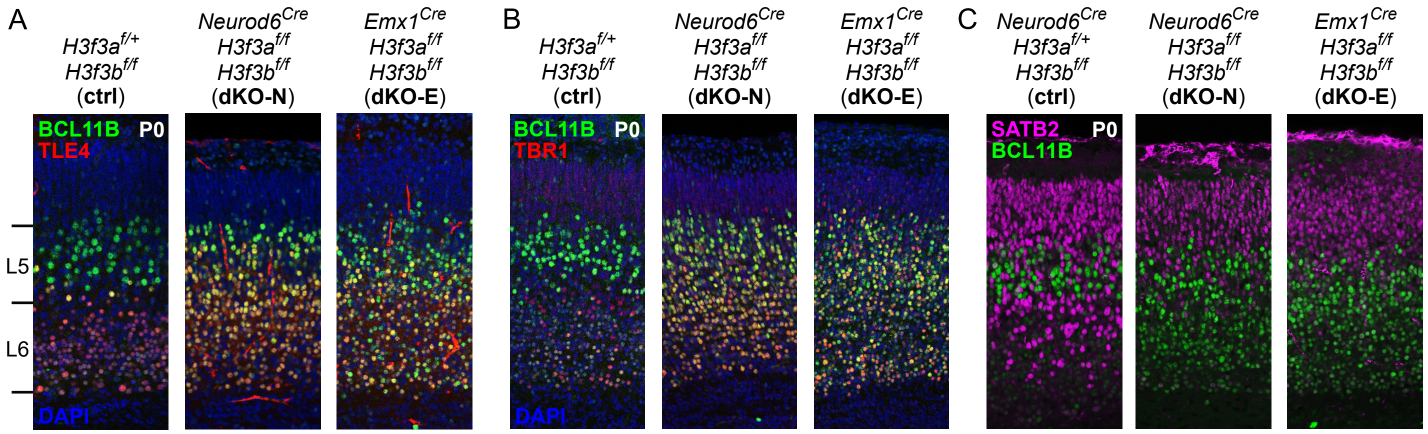


**Fig. S5**

(A and B) Coronal sections of P0 ctrl, dKO-N, and dKO-E analyzed by layer marker co-immunostaining. In control, L6 neurons were intensely labeled by TLE4 (red, A) and TBR1 (red, B) and weakly labeled by BCL11B (green), and L5 neurons showed intense BCL11B labeling and an absence of TLE4 and TBR1. In dKO-N and dKO-E, L5 and L6 neurons were abundantly co-labeled by BCL11B and TLE4 (A), and BCL11B and TBR1 (B). (C) Coronal sections of P0 ctrl, dKO-N, and dKO-E analyzed by SATB2 (magenta) and BCL11B (green) co-immunostaining. In control, SATB2 and BCL11B labeled largely non-overlapping neurons, with only a small proportion of neurons showing co-labeling. In dKO-N and dKO-E, this separation was maintained, and upper layer neurons did not misexpress BCL11B.

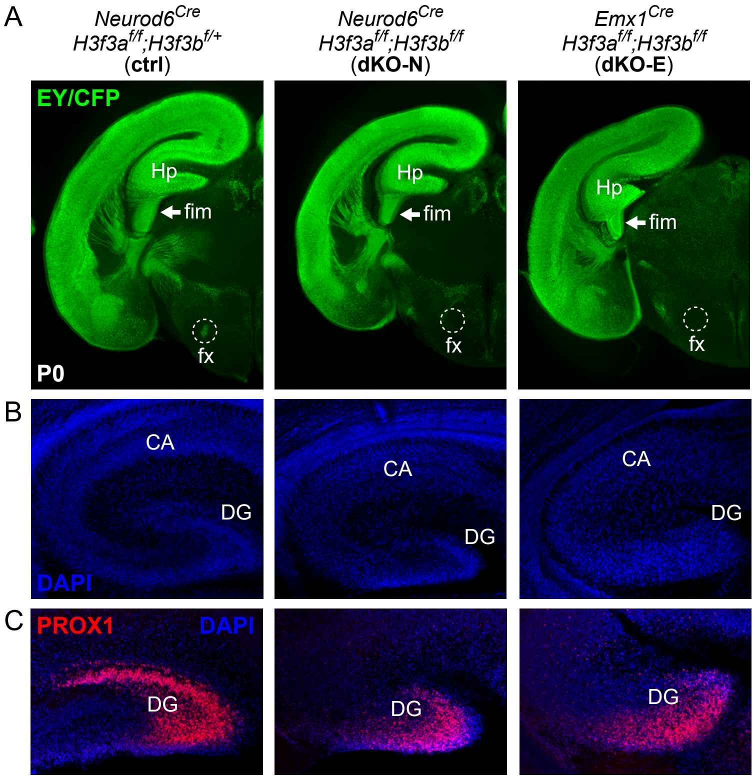


**Fig. S6**

(A) Axon analysis by EGFP immunostaining of Cre-dependent reporters (EY/CFP, green). In P0 control, EY/CFP-labeled axons from hippocampus (Hp) extended through the fimbria (fim) and innervated the fornix (fx). In dKO-N and dKO-E, hippocampal axons projected into fimbria (arrows) but failed to innervate fornix (dashed outlines). (B) DAPI staining (blue) of P0 hippocampus revealed dysplasia of cornu ammonis (CA) and dentate gyrus (DG) in dKO-N and dKO-E. (C) PROX1 immunostaining (red) of P0 hippocampus revealed disorganization of dentate gyrus cells in dKO-N and dKO-E.

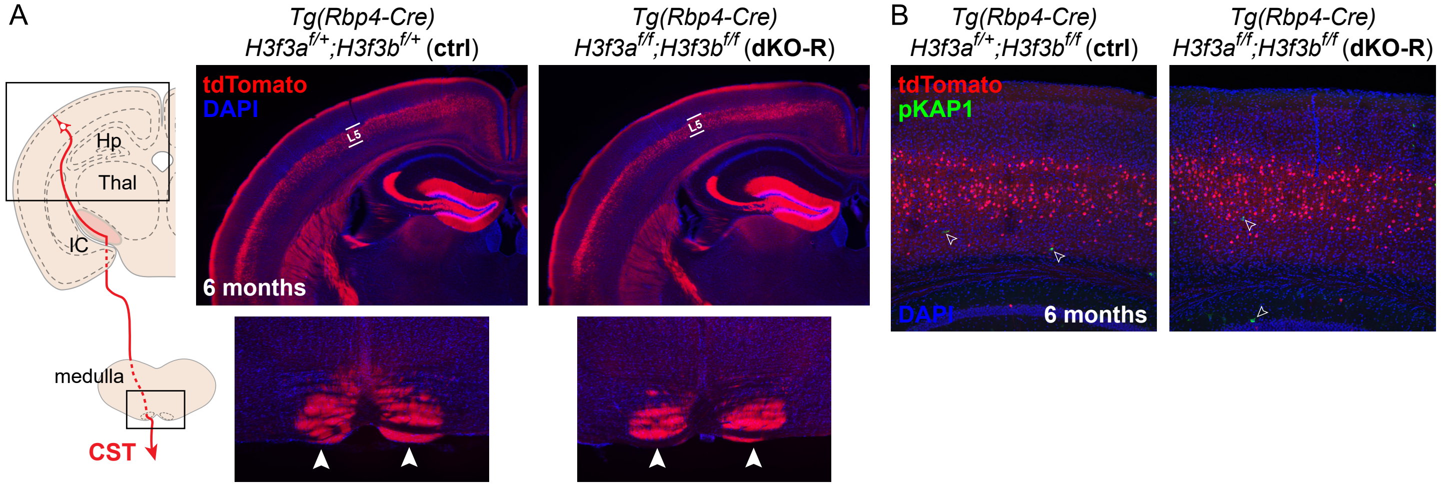


**Fig. S7**

(A) In Tg(*Rbp4-Cre*)*;H3f3a*$^{f/f}$;*H3f3b*$^{f/f}$ (dKO-R) at postnatal 6 months of age, tdTomato-labeled axons (red) arising from L5 neurons abundantly innervated the corticospinal tract (CST) and reached the pyramids (arrowheads) at the level of the medulla in a manner indistinguishable from control. (B) Immunostaining of DNA double strand break marker pKAP1 (green) in 6-month cortex. TdTomato-labeled neurons (red) in dKO-R showed no increase in pKAP1 immunostaining compared to control.
